# Mapping dendritic spines using 2D two-photon laser scanning

**DOI:** 10.1101/2025.07.10.664064

**Authors:** D. Cupolillo, V. Regio, A. Barberis

## Abstract

Neurons transform complex spatiotemporal patterns of synaptic input into structured sequences of action potentials that relay meaningful information. Excitatory inputs converging onto dendrites engage or interact with local non-linear regenerative events, adding a computational layer that expands the variety and complexity of input-output transformations. In conjunction with non-linear conductance-mediated mechanisms, the spatial arrangement of active synapses along dendrites strategically shapes this local computation. Mapping synaptic input organization is thus critical to fully uncover neuronal input-output function. Spine calcium imaging offers a direct functional readout of the location of active contacts, but such mapping requires access to spines distributed on intricate three-dimensional dendritic trees. We present a modular software pipeline for targeted imaging and analysis of dendrites using sequential two-dimensional scanning on standard two-photon microscopes. Designed to work with conventional two-photon microscopy setups, the method is fully compatible with ScanImage. It includes a pre-acquisition tool (*ROIpy*) and a post-acquisition analysis suite (*Spyne*). *ROIpy* generates dendrite-aligned region-of-interests for scattered depth-specific acquisition of neuronal arborizations. *Spyne* includes deep-learning modules for spine identification (using DeepD3) and within-spine calcium events detection (via a custom-classifier). This method is compatible with a range of experimental designs, including simultaneous two-photon imaging and patch-clamp recordings, as well as fully optical setups. The acquisition pipeline supports plane-by-plane imaging of the whole-arbor, targeted to specific compartments or to user-defined branches of interest. Our work provides a versatile strategy for targeted dendritic imaging using two-dimensional scanning multiphoton microscopy. While *ROIpy* allows adaptation to diverse experimental goals beyond synaptic mapping, *Spyne* provides an analysis strategy for functional mapping of active synapses at single-cell resolution, offering a basis for modeling how the spatial organization of synaptic inputs shapes dendritic integration..

## Introduction

The complexity of cognitive functions reflects the capacity of individual neurons to process a multitude of information and return appropriate responses [1,2]. Neurons continuously transform arrays of inputs received onto thousands of dendritic spines into meaningful neural code in the form of action potential (AP) patterns. The variety of neuronal input-output (I-O) transformation is enhanced by stacking layers of non-linear integration. This model holds that dendrites locally process inputs as an independent unit and feed the output to layers below [3–5]. For instance, in pyramidal neurons (PNs), large depolarizations in dendritic compartments produce local regenerative events such as dendritic spikes (d-spikes) through the activation of Na^+^ voltage-dependent channels (Losonczy and Magee, 2006) or slow plateau potentials through voltage-gated Ca^2+^ channels (VGCC) [6] and N-methyl-D-aspartate receptors (NMDAr) [7–10]. In combination with the distribution of active conductances, the spatial organization of inputs throughout the dendritic tree influences such local responses [8,11]. In PNs, clusters of co-active spines can trigger the generation of fast, low-traveling d-spikes (like Na^+^ spikes), producing supralinear summation of inputs and enhancing coincidence detection [12–15]. On the other hand, long-lasting plateau-like events mediated by NMDAr and VGCC support the integration of spatially or temporally dispersed input sequences across behavioral timescales [16,17]. Understanding dendritic integration within defined temporal windows requires dissecting the temporal and spatial arrangement of synaptic inputs, their identity, and their interaction. Two-photon glutamate uncaging has revealed key mechanisms of nonlinear dendritic processing and synaptic interaction [12,18,19], but it lacks information on presynaptic identity, relies on artificial input patterns, and is typically restricted to sub-portions of dendritic arbors. Anatomical labeling approaches such as GRASP (GFP reconstitution across synaptic partners) effectively mapped the whole-cell distribution of identified synaptic inputs [20–22] but do not capture temporal and functional dynamics. Calcium imaging, by contrast, provides a direct readout of synaptic activation. When combined with optogenetics, it enables inference of input identity: for example, through activation of opsins specifically expressed in defined subpopulations. For spine-level calcium imaging, laser scanning confocal or multiphoton microscopes assessing Ca²⁺-sensitive GECIs (genetically encoded calcium indicators, *e.g.*, GCaMP6s) or synthetic dyes (*e.g.*, Fluo-4FF, Fluo-5F, Fura-2, OGB-1) remains the method of choice for both *ex vivo* and *in vivo* settings [23–28]. Laser scanning strategies must balance temporal resolution, spatial coverage and spatial resolution. For instance, galvo-based systems can perform targeted scanning within spatially confined regions to increase frame rate and resolution [29]. Nevertheless, sequential acquisition of two-dimensional ROIs across multiple z-planes can fully capture the three-dimensional topology of dendritic arbors, generating a volume of discrete z layers with sparse temporal resolution. Here, we describe a modular pipeline for mapping dendritic spines activated by selected inputs using sequential 2D two-photon scanning microscopy. The pipeline is built on two main packages: *ROIpy* and *Spyne*. *ROIpy* is a pre-acquisition tool for generating 2D ROIs aligned with dendritic trajectories which integrate with scattered scanning featured by the popular data acquisition software ScanImage [30]. *Spyne* is a post-acquisition data analysis tool which includes automatic identification of dendritic spines using DeepD3 [31], followed by spine calcium event detection. For the latter, we propose a deep learning-based classifier to detect the presence of a calcium spike across thousands of spines. In this work, we report on the pipeline in its most comprehensive experimental setting, combining full dendritic calcium imaging with simultaneous patch-clamp recordings. Nonetheless, the approach offers high flexibility: it extends to all-optical configurations-combining GECIs and optogenetics-and can be tailored to target specific dendritic compartments or smaller tree subregions based on experimental needs. Overall, the approach enables efficient, high-throughput analysis of dendritic inputs across broad arbor segments without requiring volumetric scanning.

## Results

### *ROIpy* produces rectangular ROIs aligned with dendritic trajectories

We developed *ROIpy* as the pre-acquistion module of the pipeline for targeted dendritic imaging (Fig. 1A). *ROIpy* generates small, scattered 2D ROIs, each capturing short dendritic segments at discrete z positions. Iterative imaging of coplanar ROIs at successive depths allowed full reconstruction of the dendritic arbor in a layer-wise fashion. A graphical user interface (GUI, <u>Fig.</u> <u>S1A</u>) supports semi-automated ROI placement during acquisition, aligning each ROI with dendritic segments at their corresponding depths. To generate the structural reconstruction required for ROI generation, we patched ventral hippocampal CA1 pyramidal neurons (vCA1 PNs) in acute brain slices obtained from adult mice, filling them with Alexa 594 (A594) and Fluo-5F through the patch pipette to visualize dendritic morphology and monitor calcium activity, respectively. Following dye loading, we acquired a z-stack of the whole dendritic arborization (Fig. 1B) with a z step of 1-1.5 µm. We used this stack as reference to reconstruct the three-dimensional morphology of the filled neuron through the Simple-Neurite-Tracer (SNT) plug-in of ImageJ-Fiji [32], producing a standard *swc* file [33]. This format encodes neurons as a set of interconnected nodes with x, y, z coordinates, parent-child connectivity, radius, and compartment identity (Fig. 1C). *ROIpy* parses these nodes to generate rectangular ROIs and stores them into a set of *roi* files (Fig. 1B) –one per imaging plane– compatible with ScanImage’s multiple ROI (mROI) imaging tool. ROI generation is a multi-step process which includes initial ROI placement, shape adjustment and pixel allocation (see <u>Methods</u>). Shortly, nodes are first subdivided by z-plane, then by grouping uninterrupted sequences with direct parent-child connectivity. These finite dendritic segments guide initial ROI placement and shape. ROIs were then post-processed to achieve final configuration. Wider-than-longer rectangles are either eliminated or combined into a single, longer rectangle (Fig. S2B, S2C, S2D). ROIs capturing curving dendrites are extended along the width axis to ensure coverage. Then, ROIs were uniformly stretched along their major axis by a fraction of their height (see <u>Methods</u>). To avoid redundant scanning, overlapping ROIs wherein spatial overlap exceeded a defined threshold were discarded (<u>Fig. S2D</u>). The final distribution and shape of ROIs per plane depends ultimately on the density of dendritic segments crossing them. As a result, some z-layers were packed with ROIs, while others contained few. Given this scan area variability, pixel allocation was critical to control acquisition frame rate. To maintain a constant imaging rate across z-layers, we adjusted pixel dimensions independently per z-layer (Fig. 1F). Using an initial pixel dimension, we calculated the scan time per frame based on acquisition parameters—*i.e.* sampling rate, pixel dwell time, bin factor, fly-to and fly-back duration, and fill fraction (Eq. 1). We then iteratively adjusted pixel dimensions and recomputed frame rate (Eq. 2) until the desired acquisition timing was achieved. Such adjustment was constrained to the diffraction-limited resolution of the system (Eq. 3) and Nyquist sampling requirements (Eq. 4). Fine frame rate adjustments were made by modifying flyback duration, that is, the time required for the laser to return to the starting position after each frame. ScanImage stacks coplanar ROIs to minimize data volume; these are then separated during analysis to retrieve original ROI dimension. Besides the main function of ROI generation, *ROIpy* includes a suite of morphological visualization and basic structural analysis. *ROIpy* handles individual dendritic branches, *i.e.* neurites, as segments included between an initial node (the soma or a fork node) and a terminal node and can sort them by branching order or compartment type (*e.g.* apical, basal, axons) (<u>Fig. S3A</u>). To showcase the broad applicability of *ROIpy*, we analyzed a neuronal morphology dataset obtained from the Allen Cell-Types Database [34] of 63 fully-reconstructed dendritic arborizations from layer-5 (L5) PNs of primary visual areas (VISp) (<u>Fig. S3A</u>) (summarized in Table 2). Using *ROIpy*, we calculated a mean total dendritic length of 4.46 ± 0.17 mm across 47.9 ± 2.0 branches on average (<u>Fig. S3B</u>). Sholl analysis [35] revealed a branching profile consistent with L5 PNs [36], characterized by dense limited basal arborization within 200 µm from the soma and oblique apical dendrites (within 200 µm from the soma) stemming from a main trunk (elongating for up to 400 µm) that in turn furcates at the tuft beyond 400 µm (<u>Fig. S3C</u>). Convex hull analysis of terminal nodes showed a dendritic extension of 0.0085 ± 0.0006 µm^3 (^<u>Fig. S3D</u>).

**Figure 1.**
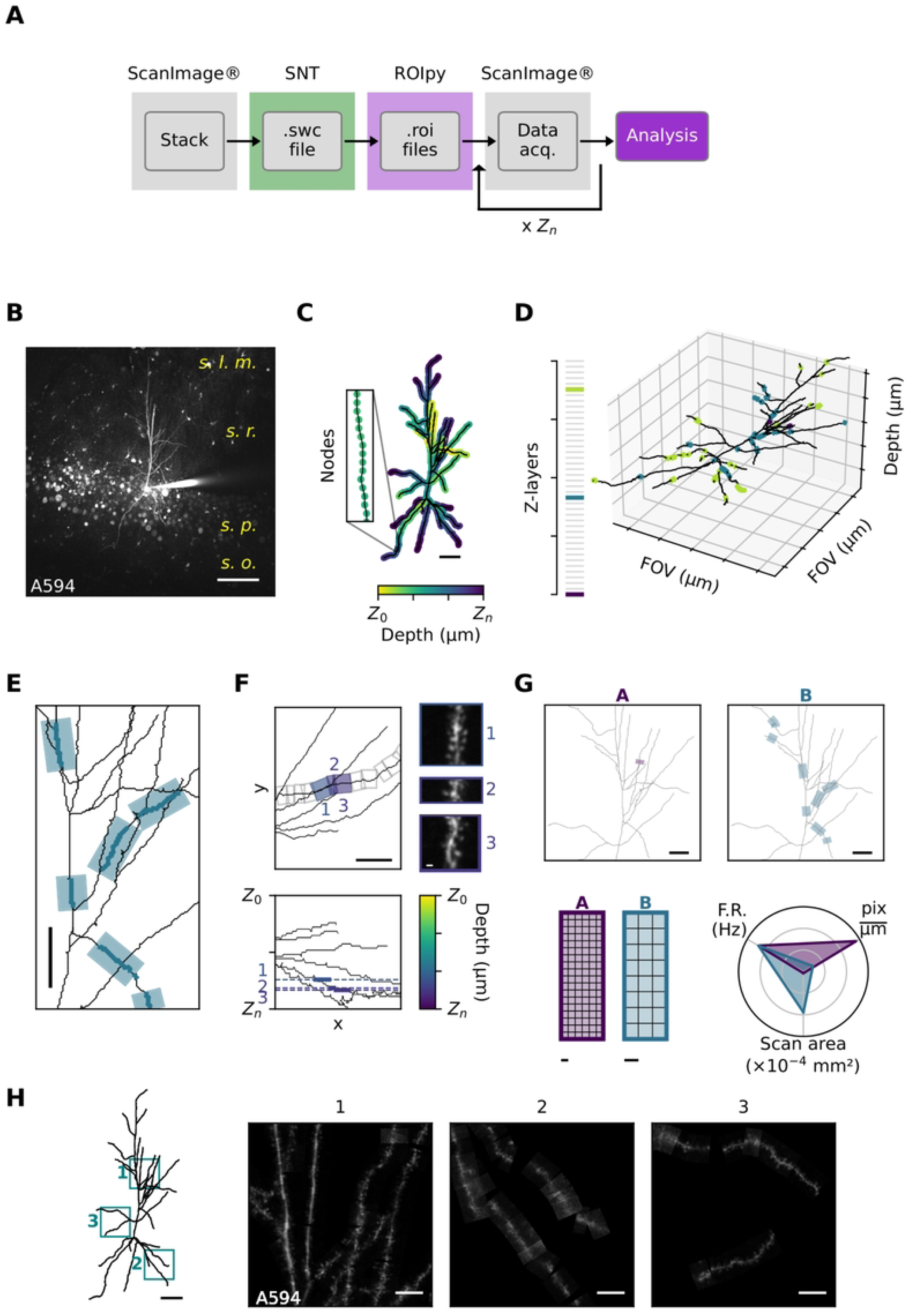
Overview of the ROI generation pipeline. **A.** Diagram of the proposed pipeline workflow. First, a reference volume is acquired using the Stack tool of ScanImage, thus ensuring that relevant metadata for subsequent ROIs placement is preserved. The volume is then used to reconstruct the neuron in 3D using tools such as the SNT plug-in in Fiji. Both the stack and the resulting. *swc* files are loaded into *ROIpy*, which generates a series of ScanImage-compatible *roi* files defining ROI placement across z-layer. Each *roi* file is sequentially loaded into the mROI module of ScanImage for targeted imaging. The acquired data are then processed using the analysis platform *Spyne*. **B.** Representative maximum intensity projection of a reference volume acquired using ScanImage. The neuron shown is a patched CA1 PN from ventral hippocampus filled with A594 and Fluo-5F. Anatomical layers are annotated: “s. o.”: *stratum oriens*, “s. p”: *stratum piramidale*, “s. r.”: *stratum radiatum*, “s. l. m.” *stratum lacunosum moleculare*. Scalebar: 100 µm. **C.** Morphological reconstruction of the neuron highlighting z-depth information for each node. Inset: zoom-in showing a segment of connected nodes. Scalebar: 50 µm. **D.** Left: three example z-layers indicated (colored) among all z-layers used to acquire the stack (gray). Right: 3D projection of the reconstructed neuron. Nodes falling within each layer are highlighted. **E.** Representative image showing coplanar ROIs placed on continuous dendritic segments within a single z-layer. Scalebar: 20 µm. **F.** Top left: Detail of ROIs placed along a dendritic branch (gray boxes), shown in 2D (x, y). Three representative ROIs are color-coded by their depth value (z). Scalebar: 20 µm. Bottom left: Side projection view (x, z) of the same dendritic tree portion, showing the z position of selected ROIs. Top right: maximum intensity projection of A594 fluorescence showing trajectory continuity of the dendritic segment despite ROIs being located on different z-layers. Scalebar: 10 µm. **G.** Top: two example z-layers from the same neuron with different ROI densities: (A) sparse ROIs, (B) more densely tiled ROIs. Bottom left: schematic showing how pixel dimensions are dynamically allocated to maintain a constant frame rate across layers. ROIs in (A) are imaged with more, smaller pixels, whereas ROIs in (B) with fewer, larger pixels. Scalebar: 1 pixel. Bottom right: radar plot illustrating the trade-off between scan area, pixel resolution, and frame rate. **H.** Left: reference volume projection of a vCA1 pyramidal neuron with three zoom-in boxes. Scale bar: 50 µm. Right: in-place ROIs highlighting the spatial distribution of dendrites and spines across the neuron. Scale bar: 10 µm.

### Spine semantic segmentation and calcium imaging

To map dendritic spines and identify those targeted by a defined axonal input, we used evoked calcium signals within spine heads as a readout of active synaptic contacts. Stimulation was delivered either electrically or optogenetically via channel-rhodopsin (ChR2). For electrical stimulation, we placed an electrode in the *stratum radiatum* of ventral hippocampal slices to stimulate the Schaffer collaterals (SCs) bundle originating from vCA3. For optogenetics stimulation, we injected CaMK2α-ChR2-EYFP into the basolateral amygdala (BLA), to stimulate projection terminals within vCA1 [37]. During scanning, we delivered either five electrical paired pulses (PPs) at 0.1 Hz or five single blue light pulses (470 nm, 2 ms) at 0.1 Hz, both evoking subthreshold excitatory post-synaptic currents (EPSCs) in vCA1 PNs. To prevent detector saturation and photodamage associated with concurrent imaging and optogenetic, we used gated photomultipliers tubes (PMTs) which allow for millisecond-timescale electrical gating. Independently of the type of stimulation, we iterated this protocol for every z-layer, recording A594 and Fluo-5F signals from *ROIpy*-generated ROIs. Data was processed using *Spyne*, a custom analysis suite for offline image inspection, stimulation artifact correction, spine segmentation, and calcium event detection. Raw data were unpacked into 2D frame sequences for each ROI. Pre-processing included 3D median filtering and correctionof optical artifacts as needed. A reference image for each ROI was generated by computing the maximum intensity projection of the A594 signal across all frames and trials, effectively revealing a high-contrast delineation of dendrites and spines (Fig. 2A). This reference image was used by DeepD3 to obtain a pixel-wise prediction map of spine probability (<u>Fig. S4D</u>) (Fernholz et al., 2024). We then transformed the probability map into a binary mask of individual spines, and applied further binary image processing (<u>Fig. S4D</u>, see <u>Methods</u>). We computed spine centroids and re-mapped them into the full FOV coordinate space using pixel-to-reference matrix transformation. In this way, we obtained the map of dendritic spines on imaged dendrites (Fig. 2B). We extracted Fluo-5F signal encased by spine masks across time (Fig. 2C) and computed Δ*F*⁄*F*_0_ by averaging pixel values and normalizing them to a rolling percentile baseline (Eq. 5). We further denoised the time series (t-series) by applying a modified Okada filter [23] (Fig. 2D). To standardize and normalize data across neurons and trials, we z-scored the traces (Eq. 6) for subsequent detection of calcium events (Fig. 2E).

**Figure 2.**
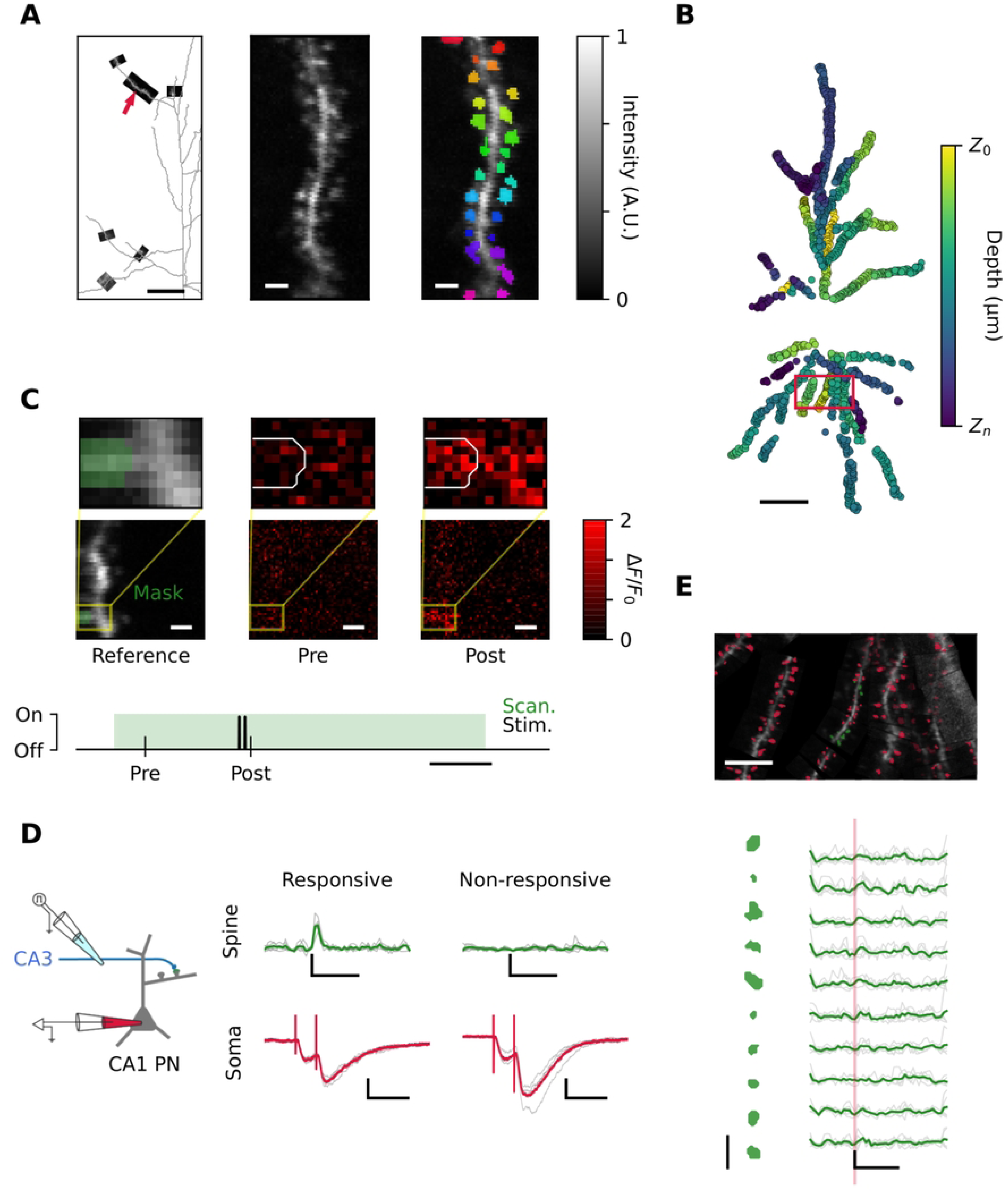
Spine semantic segmentation and calcium imaging. **A.** Left: example ROIs laying on a single z-layer aligned with dendritic morphology (gray). A representative ROI is indicated with a red arrow. Scalebar: 25 µm. Center: reference image of the indicated ROI generated by maximum intensity projection across all frames and trials. Right: pixel-wise spine segmentation masks overlayed on the reference image. Scalebar: 2 µm. **B.** Two-dimensional projection of spines centroids mapped across the whole field-of-view. Each dot marks a spine’s (x, y) position, color-coded by depth. Red box indicates zoomed view of branches shoen in E. Scalebar: 50 µm. **C.** Top: selected frames from a representative recording. Top-left: reference image with an example spine indicated in green. Top-center and top-right: frames captured before and immediately after PP stimulation, respectively, colored by ΔF/F_0_. Scalebar: 50 nm. Bottom: Digital-to-analog square waveform of the PP stimulation (red) aligned with the two-photon scan acquisition window (green). Example frames timestamps indicated. Scalebar: 500 ms. **D.** Left: schematic of the stimulation-imaging configuration. Right: representative traces from a responsive and a non-responsive spine (top) and the corresponding somatic EPSCs (bottom). Individual trials in gray and average in green (spine) and red (soma). Scalebars: calcium traces, 1 s and 0.5 ΔF/F_0_; EPSC, 100 ms and 100 pA. **E.** Top: in-place ROIs from the box highlighted in B, with spine masks in red. Example spines are indicated in green. Scalebar: 10 µm. Bottom: selected example spines (left) with corresponding calcium activity (right). Individual trials are shown in gray and average in green. The red vertical line indicates stimulation. Spine masks scalebar: 2 µm. Calcium traces scalebar: 1 s, 0.5 ΔF/F_0_ units.

### Z-score classifier

Manual detection of calcium events across thousands of spines is labor-intensive and prone to errors and inconsistencies. To streamline data processing and automate analysis, we developed a supervised binary classifier to detect calcium events in discrete, univariate z-score t-series. We trained the classifier using a ground-truth dataset of 3541 expert-annotated traces, labeled as events (1) or no-event (0). The dataset was imbalanced, reflecting the sparseness of spine activation: class 0 and class 1 represented 73% and 27% of data, respectively (ratio 3:1). Data were stratified and split into a training and a validation set (0.75/0.25, n = 2578/963 t-series respectively) and used to train a convolutional neural network (CNN) with pooling layers typically used for time series classification [38]. During training, we monitored the F1 score. Model hyperparameters, regularization and data augmentation parameters (rate and amplitude) were set using a Bayesian search method whose goal is to minimize validation loss while maximizing validation F1 score and area under the precision-recall (PR) curve (Table 4). The best-performing model converged within 24 epochs, with training and validation loss functions decreasing at comparable rates, while F1 score shows a steep rise during the initial epochs indicating a fast improvement in the balance between precision and recall, followed by a plateau that suggests convergence toward stable performance. On an independent never-seen-before test dataset made of 360 z-score t-series (class ratio 3:1), the trained model achieved an area under the PR curve of 0.93. We then established a decision boundary by selecting the probability threshold that maximized the F1 score on the test dataset to convert predicted probabilities into binary outputs (<u>Fig. S6A</u>). When evaluated against the ground truth, the classifier achieved a precision of 95.2% for predicted class 0 and 86.5% for predicted class 1, reflecting reliable classification for both classes (Fig. 3C and <u>Fig. S6A</u>). Recall was 95.5% for class 0 and 85.5% for class 1, indicating high sensitivity. To examine the internal representation learned by the network, we extracted activations from the final hidden layer (just before the sigmoid output) for all samples in the datasets and projected them into a two-dimensional space using Uniform Manifold Approximation and Projection (UMAP) [39,40]. The low-dimensional representation revealed consistent segregation of traces across datasets according to their ground-truth labels, indicating that the network learned a generalizable latent representation (<u>Fig, 3D</u>). We benchmarked performance against two standard methods: threshold-crossing and logistic regression. For the former, we tested cut-off values of 1.96 and 2.5 times the standard deviation (σ). The 1.96σ threshold showed slightly reduced accuracy (89.4%), indicating a higher rate of false positives. Similarly, the 2.5σ threshold showed an accuracy of 87.7%, but with a class-1 sensitivity of 65.6% (<u>Fig. S6B</u>). Finally, we fit the labeled training dataset to a logistic function. Logistic regression separated the test dataset based on the dot product of weights and z-score features (<u>Fig. S6C</u>). After optimizing the F1 score for thresholding, the regression model yielded slightly lower recall for 0-labeled traces (91.1 %) and 1-labeled traces (82.2 %) as compared to our classifier (<u>Fig. S6C</u>Identifying calcium events from thousands of dendritic spines acquired as previously described is a tedious and error-prone task. To streamline data processing and automate analysis, we developed a supervised binary classifier to detect calcium events in discrete, univariate z-score t-series of calcium signals. The classifier was trained on a dataset of 3541 human-annotated traces, labeled according to the presence or absence of calcium events (labeled as 1 or 0, respectively). The dataset was split into a training and a validation set (0.75 – 0.25 proportions, 2578 – 963 t-series respectively), used to train a convolutional neural network (CNN) with pooling typically used for time series classification tasks [38]. An independent never-seen-before test dataset made of 870 z-score t-series was used to evaluate the obtained model’s performance. Training hyperparameters, regularization and data augmentation rate and amplitude were set using a Bayesian search method to maximize validation accuracy (Table 3). During training, loss functions decreased and converged before reaching 20 epochs, while accuracy steeply increases and then stabilizes, achieving a max. accuracy of 97.6 %. The trained model achieved an area under the receiver operating characteristic (ROC) curve of 0.94. When ranked according to the probability of containing a calcium event, predictions well mirrored human scoring (Fig. 3C and <u>Fig. S5A</u>). To examine the internal representation learned by the network, we extracted the neuron activations from the final hidden layer (just before the sigmoid output) for all samples in the training, validation, and test datasets and projected them into a two-dimensional space using Uniform Manifold Approximation and Projection (UMAP) [39,40]. The resulting low-dimensional representation revealed consistent segregation of traces across datasets according to their ground-truth labels, indicating that the network learned a robust, generalizable mapping from input traces to a separable latent space (<u>Fig, 3 D</u>). To benchmark the model, we compared its classification performance against simpler detection approaches, specifically traditional threshold-crossing and logistic regression. For our classifier, we first established a decision boundary by selecting the probability threshold that maximized the F1 score on the test dataset, allowing us to binarize the predicted probabilities into binary outputs (Fig. S5A). When evaluated against the human-annotated ground truth, our classifier correctly predicted 99.6% of the 0-labeled z-score time series and 75.0% of the 1-labeled z-score time series (Fig. S5A). For threshold-crossing classification, we tested cut-off values of 1.96 and 2.5 times the standard deviation (σ) of each time series (Fig. S4B). Compared to the ground truth, classification based on the 1.96 × σ threshold showed slightly reduced accuracy for 0-labeled traces (98.2%, indicating a higher rate of false positives), while the 2.5 × σ threshold resulted in a markedly reduced ability to classify 1-labeled traces (46.9%, reflecting more false negatives) (Fig. S5B). Finally, we fit the labeled training dataset to a logistic function. The resulting logistic regression well separates the test dataset in the two classes, as shown by the dot product of the fitted weights and the z-score features (<u>Fig. S5C</u>). Similarly, we established a decision boundary that maximizes F1 score and used it to evaluate the model performance against the test dataset. The logistic model yielded lower sensitivity for 1-labeled traces (62.5 % of true positives), resulting in a higher false negative rate, while showing comparable performance on 0-labeled traces (99.8 % of true negative) (<u>Fig. S5C</u>). These results emphasize the advantage of our classifier in capturing calcium events that may be missed by simpler linear decision boundaries. Although logistic regression showed good accuracy (88.8%), our classifier showed an accuracy of 93.1%. We derived contingency tables to compare binary predictions from our custom classifier to those of the other classification methods, using ground-truth test labels (<u>Figure S6D</u>). Our custom classifier produced 19 more correct classifications than the 1.96σ (McNemar’s test, *p* = 0.03098) and 26 more than the 2.5σ threshold (*p* = 0.00355). Similarly, the classifier outperformed logistic regression, with 21 additional correct predictions (*p* = 0.01402). These results indicate that our custom classifier learns features that generalize better than fixed thresholds or linear approaches, especially for detecting sparse events in imbalanced data.

**Figure 3.**
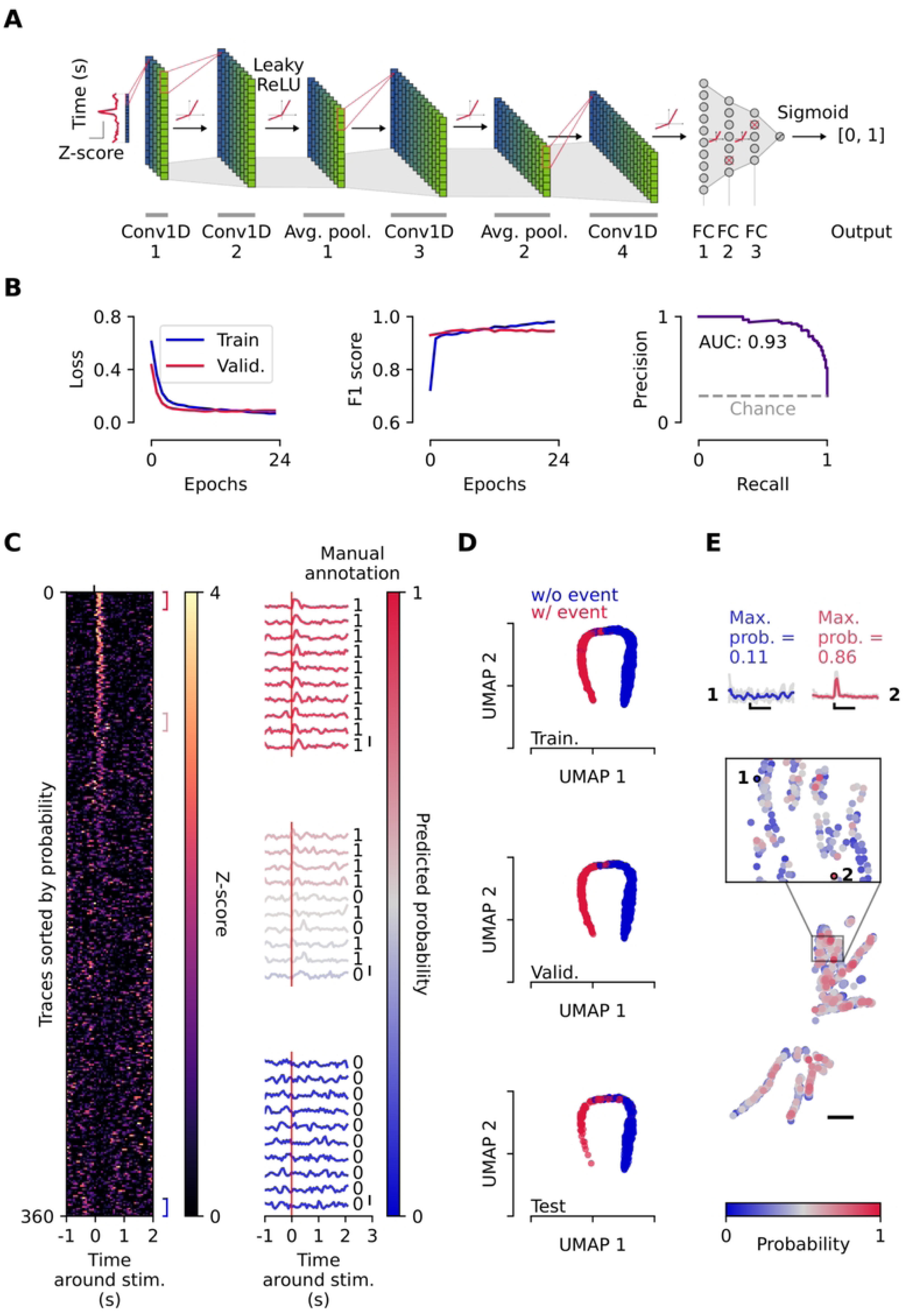
Z-score classifier architecture and performance. **A.** Simplified schematic of the neural network architecture. A 50-time points z-score trace is input to a series of 1D convolutional layers and pooling layers, followed by 3 fully connected layers. Leaky ReLU activation is applied between layers where indicated. The final output passes through a sigmoid function, yielding a probability score for event classification. The final output is thresholded and labeled [0, 1]. **B.** Training performance. Left: training and validation binary cross entropy loss decreases and converges over 24 training epochs. Center: training and validation F1 score increases and stabilizes. Right: precision-recall curve of the best performing model. The area under the curve is indicated. Gray dashed line indicates performance of a random classifier for comparison. **C.** Classifier output on held-out test dataset. Left: heatmap of test traces sorted by predicted event probability. Square brackets indicate representative ranges of high, intermediate and low probability. Right: example traces from selected low, intermediate, and high probability ranges. Ground truth labels are shown for comparison. Scale bar: 5 z-score units. **D.** Dimensionality reduction of latent features using UMAP. Top to bottom: training, validation and test dataset. Each dot represents a trace, color-coded by ground truth label. For every dataset, traces form two separable clusters, indicating effective learning features. **E.** Spatial mapping of classifier output. Bottom: dendritic spines map from a representative neuron, color-coded by the maximum event probability across 5 trials. Scalebar: 50 µm.Middle: zoom-in of a sub-portion of dendrites, highlighting two example spines. Top: z-score traces of the indicated spines. Spine 1 shows low max. event probability; spine 2 shows high max. event probability. Gray: individual trials; blue/red: mean trace. Scale bars: 1 s, 1 z-score unit.

## Discussion

Understanding how the structural organization of synaptic inputs—that is the relative spatial positioning of co-active synapses during specific behavioral contexts—shapes dendritic integration is essential to define the rules of neuronal computations [10,13,41]. Calcium transients within spine heads reliably report synaptic activity yet capturing these events across the full dendritic arbor remains technically challenging. Two-photon microscopes generate high-resolution imaging of spines on short, in-focus dendritic segments via targeted scanning [14,26,27]. Capturing extended dendritic structures, however, requires sequential plane-by-plane scanning, which is laborious, slow, and user-biased, limiting the ability to assess global spatial patterns. To overcome this barrier, we developed *ROIpy*, a Python package which, in conjunction with ScanImage mROI acquisition tool, automates the targeting of two-photon scanning to dendritic branches across depths. *ROIpy* utilizes dendritic reconstruction (encoded in the common *swc* format) obtained by any user-favorite morphological tracing software to tile two-dimensional ROIs aligned with arbor segments at the appropriate z depth. In addition, this package performs morphological analysis, enabling correlation between dendritic structure and functional synaptic map. We tested this by using an open dataset of L5 PNs reconstructions. ROI placement accuracy and dendritic coverage depends on the quality of the initial reconstruction, which in turn is contingent on the fluorescent dye–either pipette-loaded or genetically-encoded– ability to fill distal and fine branches. With 20 minutes of A594 loading, we routinely reconstructed branches 300–400 µm from the soma. Although focus-and-acquire steps require experimental time, *ROIpy* eliminates ROI preparation, reduces user bias and significantly expedites acquisition. The resulting set of acquired ROIs spans the targeted dendritic branches. This constitutes a flexible dataset wherefrom user-defined structural or functional analyses can be performed. In our case, we used ROIs to generate a static map of synaptic inputs based on a functional readout. *ROIpy* facilitates the acquisition of numerous ROIs, generating a data volume that precludes manual annotation of spine position. To address this, the post-acquisition analysis package *Spyne* incorporates DeepD3 for automated spine recognition. Existing tools for spine identification often require high-resolution 3D images for fine morphological study [42] or rely on proprietary environments (*e.g.* Matlab) [43]. In contrast, DeepD3 aligns with our needs, providing scalability and compatibility with a Python-based pipeline. We trained a custom model on our own annotated dataset of 2D images, adding a custom training hyperparameters search. Through DeepD3, we located up to thousands of spines and determined their position across branches. Traditional methods of calcium activity analysis such as peak detection, deconvolution, or thresholding are generally optimized for continuous recordings and are sensitive to baseline shifts and noise amplitude. Instead, given the structure of our data— thousands of short, discrete and normalized t-series with variable signal-to-noise ratios—we trained a custom CNN to detect stimulus-evoked events. For supervised learning, we compiled a labeled dataset of z-score traces. Manual annotation revealed a highly imbalanced class distribution, with class 0 representing over 90% of samples. To ensure sufficient representation of active synapses, we increased the prevalence of class 1 samples compared to the real distribution (5-10%) to a 1:3 ratio (963 class 1 traces instead of 150-300). To further reduce class imbalance effect, we monitored F1 score and precision-recall curve rather than accuracy throughout training and selected the model that best balanced these metrics. Model output shows an event probability distribution skewed towards 0 and 1. For this, we reasoned that an adaptive binary decision boundary should be set for each specific dataset, using F1 score or precision-recall as selection criteria, ensuring optimal tradeoff between sensitivity and precision in event detection. We also tested thresholding and logistic regression for comparison. Thresholding (using a standard variance-dependent cut-off) yielded high false positives, misclassifying noise fluctuations as events. Stricter thresholds reduced false positives but increased false negatives, overlooking small yet reliable events. Logistic regression achieved 88.8% accuracy; however, our classifier outperformed it, reaching 93.1%, with a higher recall for class 1 (85.5% vs. 82.2%) and improved precision (86.5% vs. 82.1%). Although the overall accuracy gain appears modest, our custom classifier produced 21 additional correct predictions (5.8%), a meaningful improvement in imbalanced datasets, where better calibration between signal and noise is critical. We showcased our approach using acute slice patch-clamp recordings. While this offers access to somatic subthreshold dynamics and control over resting membrane potential, it entails up to 2-3 hours of continuous recording for full arbor scanning, straining seal stability and increasing experimental failure rate. *ROIpy* is also well-suited for all-optical approaches which are less experimentally demanding. Sparse expression strategies–*e.g.* co-infection with a Cre-dependent jGCaMP and a low-titer Cre-expressing virus–enable genetically selective labeling of neurons and help resolve single-neuron morphology in dense neuropil. Future implementations may benefit from vectors co-expressing GECIs and membrane-tethered fluorophores (separated by linkers) under the control of a common promoter. In this work, we map synaptic inputs using electrical stimulation or optogenetics. To target multiple input populations, spectrally distinct opsins (*e.g.*, a blue-light sensitive ChR in combination with a red-shifted variant) can be expressed in separate afferent populations, enabling independent stimulation of multiple pathways converging on the same neuron. This requires appropriate selection of dichroic mirrors to combine and separate excitation wavelengths. Coupling optogenetic stimulation with multiphoton calcium imaging demands additional protection of the detectors. For that, we used gated PMTs to transiently disable signal collection during LED illumination. To compensate for gating-induced image loss, we applied a gating artifact correction that identifies and interpolates affected scan lines. The loss of information is negligible: we set a 6 ms gating delay after LED illumination, which falls within the range of a synaptic delay, preserving event onset. Nonetheless, in cases where the peak of the calcium transient is partially lost during gating, the slow decay kinetics of Fluo-5F [44] enable reliable detection in subsequent frames, ensuring accurate event reconstruction. We used *ex vivo* slices to report proof-of-concept for our spine mapping strategy. Slices offer precise timing of stimulation, low background activity, and facilitate optogenetic manipulation. However, *ROIpy* can be adapted *in vivo* to guide sequential targeted scanning in two-dimensional multiphoton setups. This could support functional synaptic mapping based on behavioral tasks or sensorial stimulation. For this, two extensions are required in *Spyne*: first, integration of motion correction; second, subtraction of backpropagating action potentials to unmix spine calcium signals from global calcium activity [14]. For this work, we omitted motion correction because of mechanical stability of acute slices; residual movements due to perfusion or pipette insertion are negligible over the short stimulation window (3.125 s per z-layer). Still, slow drifts accumulating during the experiment may cause misalignment between acquisition and original reference images, leading to out-of-focus ROIs and ultimately gaps in the spine map. The final output of the workflow is a functional input map with single-spines resolution. This methodology complements structural techniques such as fluorescent imaging methods [45] or electron microscopy [46] which yield dense maps but lack functional readouts. Similarly, GRASP [21,22,47] relies on interaction with specific presynaptic partners, but does not capture activity. These approaches cannot report on dynamic and activity-dependent processes under physiological or experimental conditions. Conversely, our method identifies individual spines activated by stimulation paradigms. Moreover, it permits repeated measurements for plasticity studies. While our method enables accurate mapping, resolution limits of two-photon imaging must be evaluated. Previous work comparing two-photon to beyond-light-diffraction-limit STED (stimulated emission-depletion) microscopy has shown that approximately 23% of two-photon resolved spines correspond to two distinct spines in super-resolution images [48]. As a result, some spines may reflect compound activity of closely apposed input. Despite this, the spatial and temporal resolution is sufficient to quantify input distributions across spatial scales. Furthermore, contrary to our approach, STED is not strictly compatible with high throughput functional imaging.

## Methods

### Animals

All animal care and experimental procedures were conducted in accordance with Istituto Italiano di Tecnologia (IIT) licensing as well as the Italian Ministry of Health (D.Lgs 26/2014) and EU guidelines (Directive 2010/63/EU). The experiments were conducted on Fos-TRAP2 (Fostm2.1(iCRE/ERT2) Luo/J, Jackson laboratory, USA) transgenic mice of either sex aged 5 to 7 weeks. Male and female inbred Fos-TRAP2 mice were housed in filtered cages, in a climate-controlled animal facility (22 ± 2°C) and maintained on a 12-hour light/dark cycle with water and food *ad libitum*.

### Virus injection

Mice were first weighed and checked for overall health status. Subsequently, they were placed for 3 minutes in an anesthesia chamber with isoflurane (4%) and 95% oxygen (1.5 L/min). Mice were then mounted on a stereotaxic frame under 2% isoflurane (0.8L/min). Injection of ketorolac (5-7.5 mg/kg) and enrofloxacin (5 mg/kg) were provided intraperitoneally as analgesic and antibiotic, respectively. During the surgery, the temperature was maintained at 37°C using a feedback-controlled heating pad (RWD Life Science Co., Ltd) and ophthalmic ointment was applied to avoid eye drying. A small craniotomy was performed with a dental drill (RWD Life Science Co., Ltd). AAV-CaMKII-hChR2-EYFP (titre > 10^13 vg/ul, Addgene: # 26969-AAV5) was injected bilaterally in basolateral amygdala (stereotactic coordinates relative to bregma: AP +1.6, ML: ±3.32, DV: +4.90) through a glass pipette attached to a Hamilton syringe (Model: 1701RN) at a rate of 60 nl/min controlled with a microinjection syringe pump (UMP3, World Precision Instruments, Sarasota, FL, USA). Following infusion, the injection syringe was left in place for 10 additional minutes before withdrawal. Mice were then monitored for two days post-surgery to ensure proper recovery. Mice were allowed to recover for 4 to 6 weeks.

### Acute slice preparation

Mice were deeply anesthetized by injecting intraperitoneally ketamine (200mg/kg) and xylazine (20mg/kg) and transcardially perfused with an ice-cold N-methyl-D-glucamine (NMDG)-based solution containing (in mM): 92 NMDG, 2.5 KCl, 1.25 NaH_2_PO_4_, 30 NaHCO_3_, 20 HEPES, 25 glucose, 2 thiourea, 5 Na-ascorbate, 3 Na-pyruvate, 0.5 CaCl_2_·2H_2_O, and 10 MgSO_4_·7H_2_O, 300 mOsm/Kg, pH adjusted to 7.3-7.4 with 5M hydrochloric acid and equilibrated with 95% O_2_ and 5% CO_2_. Brains were extracted and submerged in ice-cold NMDG-cutting solution. Horizontal brain slices (350 µm thick) were prepared from either hemisphere in ice-cold NMDG-cutting solution using a vibratome (VT1200S, Leica Microsystems), incubated in a maintenance chamber at ∼35 °C for 20/30 minutes in the NMDG-cutting solution, and subsequently stored at room temperature (RT) in artificial cerebrospinal fluid (ACSF) until recording. The ACSF solution contained (in mM): 125 NaCl, 26 NaHCO_3_, 2.5 KCl, 1.25 NaH_2_PO_4_, 10 glucose, 2.3 CaCl_2_, and 1.3 MgCl_2_, equilibrated at 95% O_2_ and 5% CO_2_ for at least 40 minutes.

### Patch clamp electrophysiology

Patch clamp experiments were performed similarly to previous reports [49]. After at least 1h of recovery in ACSF, slices were transferred to a recording chamber and constantly perfused with a zero-Mg^2+^ ACSF (2ml/min) at RT, to allow for the removal of the Mg^2+^ block of NMDA receptors. Picrotoxin (50 µM), d-serine (10 µM), and CNQX (10 µM) were present in the superfusate of all experiments to ensure that Ca^2+^ events originated from monosynaptic glutamate transmission. Patch pipettes were prepared using a horizontal puller (P-1000 Next Generation Micropipette Puller, Sutter Instrument) from borosilicate glass capillaries (Warner Instruments, LLC, Hamden, USA). When filled with a K-gluconate-based intracellular solution, pipette resistance was 4 to 6 MΩ. The composition of the intracellular solution was as follows (in mM): 135 K-gluconate, 5 KCl, 2NaCl, 10 Na2P-Cratine, 0.1 EGTA, 10 Hepes, 5 Mg-ATP, 0.4 Na-GTP (285 mOsm/Kg, adjusted to pH 7.2 with KOH). In addition, Alexa-Fluor 594 (75 µM, ThermoFisher #A10438) and the synthetic calcium indicator Fluo-5F pentapotassium salt (300 µM, ThermoFisher #F14221) were added to the intracellular solution. Cells were identified by a Dodt Gradient Contrast (DGC) microscope (Hyperscope, Scientifica, UK) in bright-field using a UV filter paired with an oil-condenser (Olympus U-AAc), using a 16X water-immersion objective lens (Nikon). Whole-cell configuration was reached in identified and selected pyramidal neurons in ventral CA1. Holding current and access resistance were continuously monitored throughout the experiment. Cells were discarded if holding current was > 200 pA and if access resistance > 20 MΩ or if these parameters changed by more than 25% during recording. No liquid-junction potential correction was used. Currents were sampled at 10 KHz using the 700B Axopatch amplifier (Molecular Devices, Sunnyvale, CA) and low-pass filtered at 1 kHz. For SCs stimulation, an Ag/AgCl electrode was introduced in an ACSF-filled glass pipette and positioned approximately 300 µm from the patched cell in the *stratum radiatum*. To elicit a synaptic response, we delivered squared paired pulses (PPs) at 20 Hz, consisting of two stimuli (0.8 ms duration, separated by 50 ms) to minimize synaptic failures probability. The intensity of stimulation was adjusted by using an isolated constant-current stimulator (Digitimer Ltd.) to elicit a minimal synaptic response. For optogenetics, single 2 ms-long LED pulse at 470 nm (CoolLED’s pE-300^white^) with an intensity of 3 to 7 mW (measured below the objective lens) were delivered and band-passed through a GFP filter (SCI-49002s, Scientifica, UK).

### Two-photon scanning system

The optic path consists of an excitation source provided by a femtosecond Ti:Sapphire pulsed laser (Chamaleon Ultra II, Coherent Inc.), a modulatory Pockel’s cells (Conoptics, Dambury CT), beam expander, mechanical shutter, and HyperScope (Scientifica UK) Scan Path containing tube lens, beam combiner. The two-photon microscope (SliceScope, Scientifica UK) was equipped with a dual-axis galvo-galvo module (HyperScope Y-galvanometer coupled with an additional HyperScope Resonant X-Module) and GaAsP photomultipliers (Hamamatsu). The laser was tuned to 830 nm to simultaneously excite A594 and Fluo-5F, with a power of 6–11 mW measured under the objective. This power was kept constant throughout the imaging depth. Simultaneous two-photon excitation and the LED optogenetic stimulation were enabled by a custom primary dichroic mirror (ZT590-180dcrb, Chroma Technology Corp.). Emission signals were separated using a 580 nm dichroic mirror and detected through bandpass filters centered at 525/50 nm (Fluo-5F) and 620/60 nm (A594). Frames were acquired through a water-immersion objective lens (Nikon 16x / 0.8 NA / 3 mm WD) using ScanImage software (MBF Bioscience, Williston, VT). The signal was digitized in pixels with intensity values as signed 16-bit integers (-32,768 to 32,767).

### Reference volume acquisition and morphological reconstruction

After obtaining a stable whole-cell configuration, cells were allowed to load with A594-containing intracellular solution for at least 20 minutes to ensure adequate filling of dendritic processes, including thin and distal branches. The depth range containing the whole dendritic arborization was measured and, using the ScanImage Stack Acquisition tool, a bounded z-stack was configured. The number of linearly spaced slices was adjusted to yield a z-step of 1 to 1.5 µm, balancing the trade-off between axial resolution and total number of slices. Frames (1024 x 1024 pixels) were acquired at 0.27 Hz without averaging. The stack image was then used as reference to semi automatically trace the neuronal morphology using SNT. A single point soma was created, and then the traced dendrites were manually tagged as apical or basal.

### ROI design and optimization

*ROIpy* parses *swc*-written 3D neuronal morphologies into graph representations of interconnected nodes (with associated metadata). Coplanar nodes were grouped based on continuous connectivity, forming scattered chains which served as the basis for automated placement of rectangular ROIs. For each node chain, the straight line connecting the nodes closest to and farthest from the soma defined the longitudinal axis of the rectangular ROIs, determining its height and rotation. Rectangles were symmetrically expanded to a fixed width of 10 µm perpendicular this axis, ensuring the dendrite remained centered along it. ROIs were then refined as described previously. All ROIs processing parameters are tunable and are summarized in Table 1. ROIs with extreme aspect ratios (width-to-height > 3.5) were excluded unless they were contiguous with other ROIs. ROIs were symmetrically elongated by 33% of their height. To identify spatially overlapping ROIs, we created a pairwise overlap matrix for every z-layer. ROIs with more than 90% of their area covered by another ROI were discarded. A dual-radius exclusion zone was applied to filter out dendritic ROIs located within 20 µm (basal) or 50 µm (apical) around the soma. To allocate the appropriate number of pixels per ROI, a target frame rate value was assigned (*e.g.* 16 Hz) and the corresponding scan period was computed as the inverse of the frame rate. We calculated the active scanning time by subtracting the fly-to periods between ROIs and the frame-flyback to scan start position period, both initially set at 1 ms. The total active scan area was calculated by summing the area of all ROI, corrected by the fill fraction (0.9) to account for overscan required for bidirectional scanning. Using a predefined pixel dwell time (*t*_*dwell*_) of 3.2 µs, we computed an initial pixel dimension as:

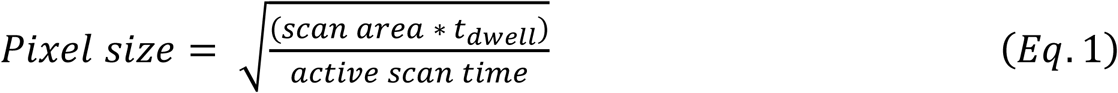

**Table 1.**

| Parameter | Value | Notes |
| --- | --- | --- |
| Initial ROI width | 10 $\mu\text{m}$ | Applied perpendicular to the dendrite axis |
| ROI elongation factor | $1.33 \times \text{height}$ | Applied along longitudinal axis |
| Aspect ratio threshold | 3.5 (width/height) | Exclusion criterion based on aspect ratio |
| Overlap removal threshold | > 90% area overlap | Exclusion criterion to avoid redundant scanning |
| Soma exclusion radius | 50 $\mu\text{m}$ (apical),<br>20 $\mu\text{m}$ (basal) | Exclusion criteria |
| Target frame rate | 16 Hz | Defines the scan period<br>(1 / frame rate) |
| Fly-to time<br>($t_{\text{fly-to}}$, per transition) | 1 ms | Non-active scanning period<br>between ROIs |
| Flyback time<br>($t_{\text{flyback}}$, per frame) | 1 ms | Return time to the starting<br>point of the frame scan |
| Fill fraction | 0.9 | Fraction of scanned area<br>accounting for overscan |
| Sampling rate | 1.250 MHz | ADC sample rate collection |
| Pixel bin factor | 4 | Number of ADC samples that<br>are averaged into one pixel. |
| Pixel dwell time ($t_{\text{dwell}}$) | 3.2 $\mu\text{s}$ | Time to acquire a single pixel |
| Max iterations | 100 | Iterative loop limit to<br>converge on target frame rate |
| Frame rate tolerance | $\pm 0.3 \text{ Hz}$ | Acceptable deviation from<br>the target frame rate |

The frame rate that resulted from such pixel dimension was calculated and compared to the target frame rate. Pixel size was iteratively refined in a convergence loop (up to 100 iterations), to achieve the target frame rate (with a tolerance of ± 0.3 Hz). To calculate frame rate, we multiplied the allocated number of pixels by *t*_*dwell*_ for each ROI, added non-active scan times:

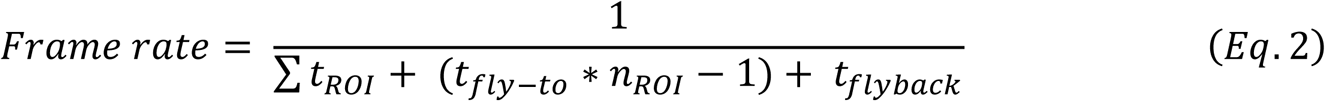

and finally ensured that the computed pixel dimensions remained compatible with both the digitizer’s acquisition rate and the two-photon sampling rate. This correction provided the actual scan duration per line, which was used to validate that the total scan time aligned with the desired frame period. Pixel dimensions were prevented from exceeding the diffraction-limited resolution of the optical instrument by defining an upper limit according to Abbe-Rayleigh’s criteria [50]:

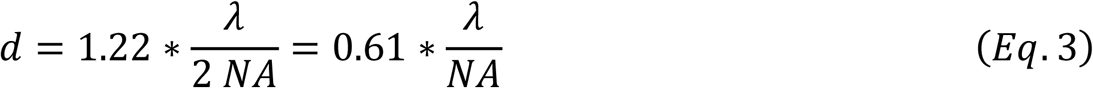

The minimum resolvable distance (*d*) between two objects depends on the numerical aperture *NA* of the microscope objective and the imaging wavelength *λ*. In line with Nyquist sampling criteria, the pixel size (*f*_*s*_) needs to be 2-3 times smaller than the microscope resolution (*f*_*max*_):

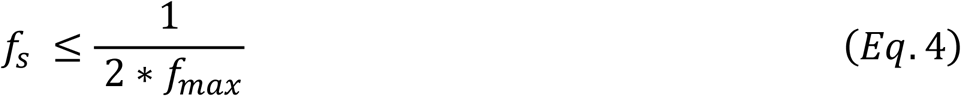

therefore, if pixel-to-µm ratio exceed this constraint during pixel dimension adjustments, ROIs are split into two or more separate scanning groups (and consequently *roi* files) located within the same z-layer, thus reducing the total scanning area per group.

### Allen cell-type dataset

We queried the Allen Cell-Type Database (https://celltypes.brain-map.org/) using an application programming interface (API) written in Python, using the *allensdk* library, to retrieve mouse neurons with available morphological reconstructions. The search was restricted to layer 5 neurons in the primary visual area (VISp). A total of 63 neurons were retrieved, and their corresponding *swc* files were used for *ROIpy*-based processing and reconstruction. Table 2 summarizes the retrieved neurons.

**Table 2.**

| **Specimen ID** | **Donor Name** | **Cre Line** | **Structure** | **Hemisphere** |
| --- | --- | --- | --- | --- |
| 485909730 | 205530 | Cux2-CreERT2 | VISp, L5 | Right |
| 323865917 | 172530 | Scnn1a-Tg3-Cre | VISp, L5 | Left |
| 314804042 | 168053 | Rorb-IRES2-Cre | VISp, L5 | Left |
| 488679042 | 212872 | Rorb-IRES2-Cre | VISp, L5 | Right |
| 560678143 | 285657 | Vip-IRES-Cre | VISp, L5 | Right |
| 471129934 | 180747 | Rbp4-Cre_KL100 | VISp, L5 | Left |
| 524850271 | 250843 | Chrna2-Cre_OE25 | VISp, L5 | Left |
| 486025194 | 205726 | Rbp4-Cre_KL100 | VISp, L5 | Left |
| 559387643 | 281934 | Ctgf-T2A-dgCre | VISp, L5 | Left |
| 480114344 | 197332 | Rorb-IRES2-Cre | VISp, L5 | Right |
| 589760138 | 318179 | Sim1-Cre_KJ18 | VISp, L5 | Left |
| 488697163 | 212873 | Rorb-IRES2-Cre | VISp, L5 | Right |
| 485574721 | 203498 | Rbp4-Cre_KL100 | VISp, L5 | Left |
| 468120757 | 177297 | Scnn1a-Tg3-Cre | VISp, L5 | Right |
| 515249852 | 243055 | Rbp4-Cre_KL100 | VISp, L5 | Left |
| 480122859 | 197352 | Rorb-IRES2-Cre | VISp, L5 | Left |
| 502366405 | 230665 | Ctgf-T2A-dgCre | VISp, L5 | Left |
| 486111903 | 205727 | Rbp4-Cre_KL100 | VISp, L5 | Right |
| 480116737 | 197332 | Rorb-IRES2-Cre | VISp, L5 | Right |
| 488698341 | 212873 | Rorb-IRES2-Cre | VISp, L5 | Right |
| 588402092 | 318850 | Tlx3-Cre_PL56 | VISp, L5 | Right |
| 480351780 | 197851 | Rorb-IRES2-Cre | VISp, L5 | Left |
| 563226105 | 291923 | Esr2-IRES2-Cre | VISp, L5 | Left |
| 607124114 | 338859 | Tlx3-Cre_PL56 | VISp, L5 | Left |
| 314822529 | 168053 | Rorb-IRES2-Cre | VISp, L5 | Left |
| 471758398 | 180748 | Rbp4-Cre_KL100 | VISp, L5 | Left |
| 471767045 | 180748 | Rbp4-Cre_KL100 | VISp, L5 | Right |
| 471143169 | 180747 | Rbp4-Cre_KL100 | VISp, L5 | Right |
| 589442285 | 318531 | Sim1-Cre_KJ18 | VISp, L5 | Left |
| 473601979 | 181715 | Rorb-IRES2-Cre | VISp, L5 | Left |
| 487664663 | 211531 | Rbp4-Cre_KL100 | VISp, L5 | Left |
| 476263004 | 187854 | Scnn1a-Tg3-Cre | VISp, L5 | Right |
| 479225052 | 187416 | Rorb-IRES2-Cre | VISp, L5 | Left |
| 480169178 | 197353 | Rorb-IRES2-Cre | VISp, L5 | Right |
| 485911892 | 204780 | Ntsr1-Cre_GN220 | VISp, L5 | Left |
| 488695444 | 212873 | Rorb-IRES2-Cre | VISp, L5 | Left |
| 485838981 | 204744 | Rbp4-Cre_KL100 | VISp, L5 | Right |
| 488680917 | 212872 | Rorb-IRES2-Cre | VISp, L5 | Right |
| 479010903 | 195278 | Pvalb-IRES-Cre | VISp, L5 | Left |
| 479225080 | 187416 | Rorb-IRES2-Cre | VISp, L5 | Left |
| 485574832 | 203498 | Rbp4-Cre_KL100 | VISp, L5 | Right |
| 478586295 | 194322 | Rorb-IRES2-Cre | VISp, L5 | Right |
| 476049169 | 187416 | Rorb-IRES2-Cre | VISp, L5 | Right |
| 479721491 | 196624 | Rbp4-Cre_KL100 | VISp, L5 | Right |
| 480003970 | 196752 | Scnn1a-Tg2-Cre | VISp, L5 | Right |
| 599334696 | 329505 | Tlx3-Cre_PL56 | VISp, L5 | Left |
| 486110216 | 205727 | Rbp4-Cre_KL100 | VISp, L5 | Right |
| 557252022 | 281330 | Sim1-Cre_KJ18 | VISp, L5 | Right |
| 354190013 | 176962 | Scnn1a-Tg2-Cre | VISp, L5 | Left |
| 515524026 | 243058 | Rbp4-Cre_KL100 | VISp, L5 | Right |
| 486052980 | 205726 | Rbp4-Cre_KL100 | VISp, L5 | Right |
| 478888052 | 193643 | Scnn1a-Tg2-Cre | VISp, L5 | Right |
| 488683425 | 203503 | Rbp4-Cre_KL100 | VISp, L5 | Left |
| 485880739 | 204745 | Rbp4-Cre_KL100 | VISp, L5 | Left |
| 471141261 | 180747 | Rbp4-Cre_KL100 | VISp, L5 | Left |
| 485835016 | 204744 | Rbp4-Cre_KL100 | VISp, L5 | Right |
| 314831019 | 168093 | Scnn1a-Tg3-Cre | VISp, L5 | Left |
| 490916882 | 215300 | Rbp4-Cre_KL100 | VISp, L5 | Right |
| 484559000 | 201965 | Rorb-IRES2-Cre | VISp, L5 | Right |
| 485837504 | 204764 | Rorb-IRES2-Cre | VISp, L5 | Right |
| 476048909 | 187416 | Rorb-IRES2-Cre | VISp, L5 | Left |
| 485835055 | 204744 | Rbp4-Cre_KL100 | VISp, L5 | Left |
| 480353286 | 197851 | Rorb-IRES2-Cre | VISp, L5 | Right |

### DeepD3 training and inference

We trained the segmentation model using a dataset of two-dimensional ROIs using the provided DeepD3 suite for training own data [31]. The dataset consisted of 27 images of dendrites and spines (min. dimension 128 x 128 pixels) characterized by an average pixel-to-µm ratio of 0.035 ± 0.010 µm, aligning with the range of the collected data (<u>Fig. S4B</u>). Images were manually annotated with pixel precision using a custom *Napari*-based GUI written in Python to generate binary masks of spines and dendrites (<u>Fig. S4A</u>). To promote detection of perpendicularly protruding spines, we labeled bright, bulbous features along the dendritic shaft as spines. 2D images and corresponding annotations were adapted to DeepD3 data streamer by introducing them in a dummy 3D stack (x, y, 1). Image stacks were preprocessed by subtracting the minimal pixel value from each frame and converted to unsigned 16-bit integers. Data was split into training and validation sets (0.8 / 0.2). Training was performed in line with DeepD3 provided toolbox. The model was trained for 30 epochs and was evaluated using a combined Dice and Mean Squared Error loss while monitoring accuracy and Intersection-over-Union (IoU). To optimize convergence, we employed an exponential learning rate decay after the 15th epoch. We customized training by adding hyperparameter tuning, performed through a Bayesian search that probed initial learning rate, number of filters in the first convolutional layer, data batching size and a minimal content threshold, using validation set accuracy as optimization target. The best performing model was chosen qualitatively based on validation accuracy, stability and convergence of training-validation loss curves and intersection-over-union (IoU) across epochs. The optimized hyperparameters are summarized in Table 3. For bulk inference, images were padded to a common size, defining height and width as the smallest multiple of 32 pixels that exceeded the largest image dimensions in the dataset. The spine and dendrites probability maps resulting from inference were then post-processed to generate final segmentations masks. Maps were binarized using a threshold of 0.3 and 0.7 for spines and dendrites respectively and spine labels smaller than 3 pixels were discarded. To eliminate segmentation artifacts disconnected from dendrites, a 12-fold dilation of the dendritic mask was applied and spine labels falling outside this boundary were removed. Additionally, to prevent artificial merging of spine masks at diagonal junctions, 2×2 windows with diagonally opposed non-zero pixels were set as 0. To separate closely apposed spines, a distance transform followed by a marker-based watershed algorithm using local maxima was applied. The resulting masks were labeled, and pixel area and centroid coordinates were calculated for each spine. Centroids were projected to FOV reference space coordinates using affine transformations.

**Table 3.**
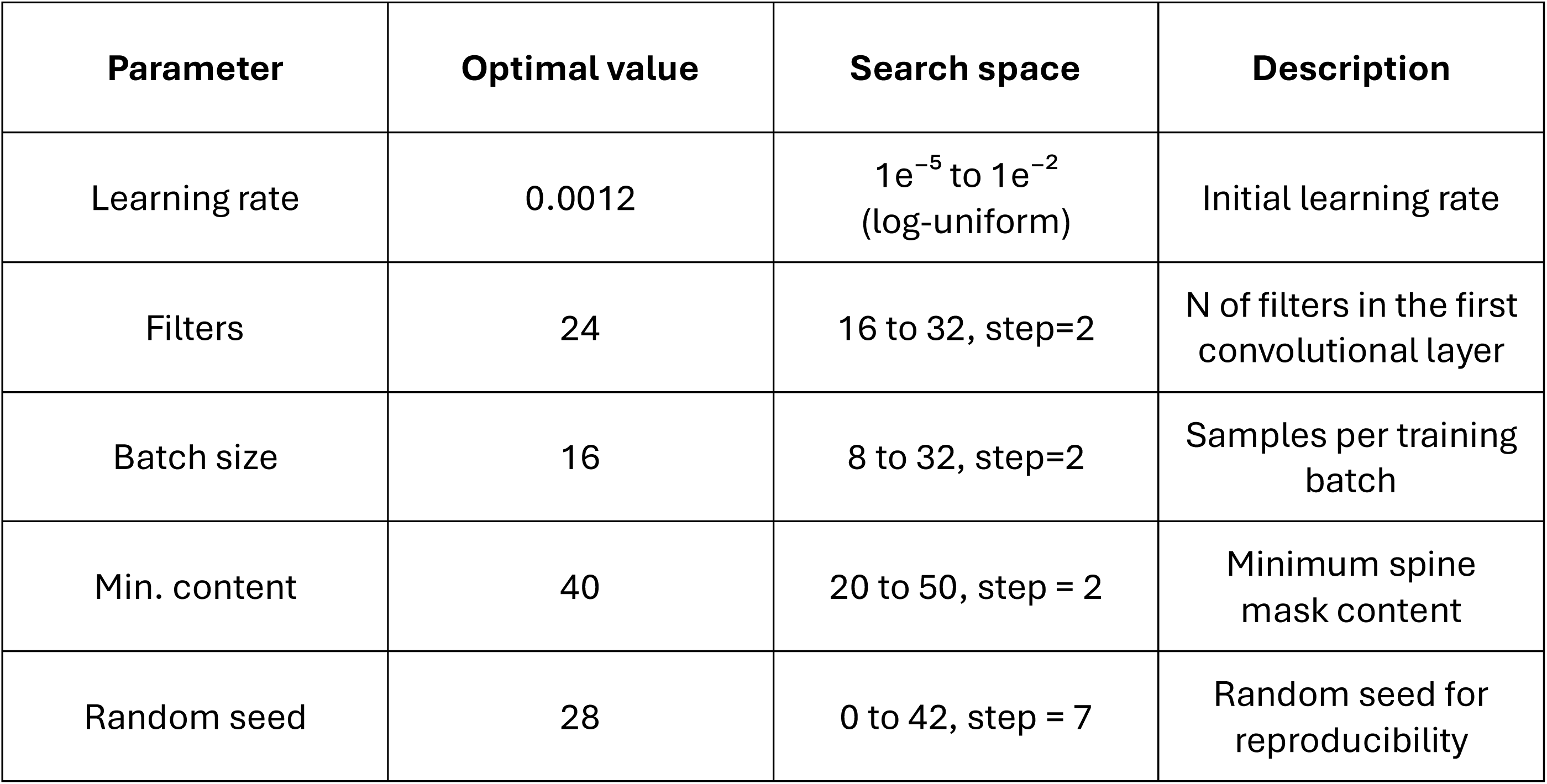

| Parameter | Optimal value | Search space | Description |
| --- | --- | --- | --- |
| Learning rate | 0.0012 | $1e^{-5}$ to $1e^{-2}$<br>(log-uniform) | Initial learning rate |
| Filters | 24 | 16 to 32, step=2 | N of filters in the first<br>convolutional layer |
| Batch size | 16 | 8 to 32, step=2 | Samples per training<br>batch |
| Min. content | 40 | 20 to 50, step = 2 | Minimum spine<br>mask content |
| Random seed | 28 | 0 to 42, step = 7 | Random seed for<br>reproducibility |

### Gated PMTs artifact correction

During optogenetic stimulation, scattering blue light used to excite ChR2 is collected by the PMTs alongside fluorescence from excited functional indicators (Fluo-5F and A-594). To protect from photodamage, PMTs equipped with gating function are employed. The gate circuit temporarily disables photon detection, electrically protecting the PMTs from excessive light exposure. The gate circuit is triggered by the Digidata 3 ms before LED onset and has a duration of 10 ms—longer than the 2 ms-long light pulse—usually within the acquisition time of a single frame of a ROI (<u>Fig.</u> <u>S5A</u>). Due to bidirectional scanning, the gating interval results in a horizontal stripe of low-intensity pixels acquired during the gating period (<u>Fig. S5C</u>). Gating artifact correction is performed as a pre-processing step. In few words, for each frame in the Alexa Fluor channel, the average intensity value of every line (row) is computed (<u>Fig. S5C</u>). Lines affected by gating are identified as those whose average intensity deviates by more than 3 standard deviations (*σ*) from the mean intensity of all lines of that frame (<u>Fig. S5C</u>). Deviating lines are then corrected by interpolating each pixel values with the average of the corresponding pixels from preceding and subsequent frame.

### Calcium imaging analysis

Frames (n = 50), which consisted of a set of coplanar ROIs, were acquired at 16 Hz, for a total duration of 3.125 s and functional signals were extracted from spine masks. A raw t-series was generated by averaging intensity of all pixels included in the mask. Δ*F*⁄*F*_0_ was calculated as:

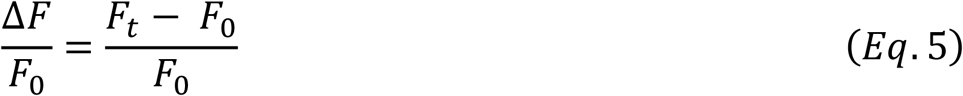

where *F*_*t*_ is the average fluorescence intensity at time *t* and *F*_0_ is the baseline trace computed as the 10^th^ percentile of *F*_*t*_ within a running kernel of 0.5 s centered in *t*. For each trace, we calculated standardized z-scores as:

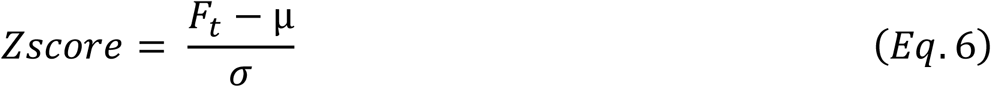

where µ is the average of all *F*_*t*_ and *σ* is the standard deviation of *F*.

### Z-score classifier training and inference

To train a classifier to identify active synapses, we built a ground-truth dataset using 3541 z-score traces randomly subsampled from recordings across various neurons. To add variability and increase model robustness, extracellular stimulation was delivered using either electrical pulses or optogenetic blue light pulses. Regardless of stimulation type, pulses were triggered 1 second after scan onset, generating time-locked calcium peaks. Traces were annotated as 0 or 1 by experts using a custom GUI written in Python. Spine activation sparseness was reflected as an imbalance towards class 0 (96.3 %) in the sampled data. Data was divided into training and validation sets (0.75 / 0.25) maintaining class ratio in both splits. The classifier was conceptualized around common CNN and pooling architecture designed for binary classification task of one-dimensional input data. The code for training was written in Python, using the deep learning library *PyTorch*. The neural network is composed by 4 convolutional layers, 2 pooling layers and 3 fully connected (FC) layers, with leaky rectifier linear unit (ReLU) activation functions. The final layer is passed to a sigmoid activation function to yield a probability score. The network was trained using a binary cross-entropy loss function between predicted probabilities and labels. Training was performed with early stopping to terminate training when the validation loss failed to improve for 10 consecutive epochs. In addition, we implemented a learning rate scheduler that reduced the learning rate by a factor of 0.5 when validation loss plateaued for 5 epochs. During training, model performance was monitored at each epoch using the F1 score:

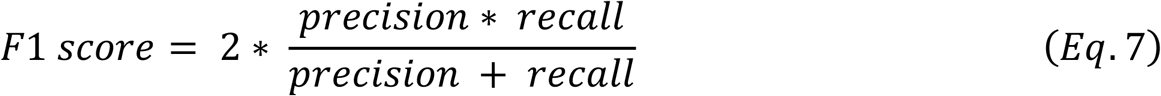

where precision and recall are:

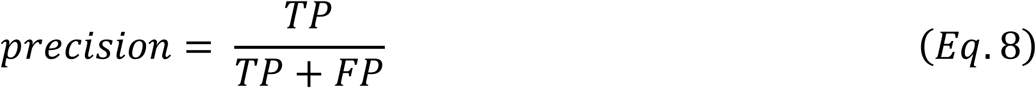

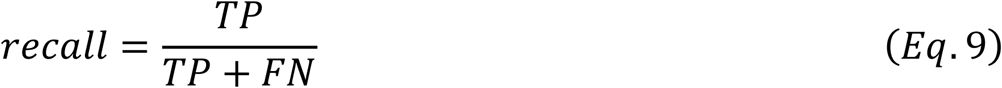

Here, TP (true positives) and TN (true negatives) are correct predictions, while FP (false positives) and FN (false negatives) represent misclassifications. To optimize model architecture and training, we tuned hyperparameters using a Bayesian search method which combined architectural parameters and training-related parameters. Architectural specifications included the number of output channels and kernel sizes for every convolutional layer, the number of units in the fully connected layers, the type of pooling operation (max or average) and the negative slope of the leaky ReLU. On the other hand, training-related parameters consisted of dropout rate, initial learning rate, optimizer settings, data batching size, random seeding, data augmentation probability (for training data only) and its relative Gaussian noise amplitude. Bayesian optimization was performed over 30 training trials, with each trial evaluated using minimization of validation loss and maximization of validation F1 score and area under the PR curve as the optimization objective. The best-performing hyperparameter configuration was then selected qualitatively based on validation F1 and stability and convergence of training-validation loss curves across epochs. The specific optimized hyperparameters and their corresponding search space are summarized in Table 4. The test dataset consisted of 360 held-out z-score traces, with a class ratio of 3:1. In all cases, the decision boundary for predicted probability values was selected by computing the F1 score across a range of thresholds and selecting the value that maximized it.

**Table 4.**

| Parameter | Optimal value | Search space | Description |
| --- | --- | --- | --- |
| Learning rate | $2.94 \times 10^{-5}$ | $1e^{-5}$ to $1e^{-2}$<br>(log-uniform) | Initial learning rate |
| L1 regularization | $1.42 \times 10^{-5}$ | $1e^{-5}$ to $1e^{-2}$<br>(log-uniform) | L1 penalty coefficient |
| L2 regularization | $0.62 \times 10^{-3}$ | $1e^{-6}$ to $1e^{-3}$<br>(log-uniform) | L2 penalty coefficient |
| Dropout rate | 0.099 | 0.0 to 0.5 | Dropout in FC layers |
| Augmentation probability | 0.086 | 0.0 to 0.5 | Probability of augmentation |
| Noise level | 0.23 | 0.0 to 0.5 | Amplitude of augmentation noise |
| Conv1 output channels | 24 | [8, 16, 24, 32] | Filters in 1st convolutional layer |
| Conv2 output channels | 32 | [16, 32, 48, 64] | Filters in 2nd convolutional layer |
| Conv3 output channels | 32 | [32, 48, 64, 80, 96] | Filters in 3rd convolutional layer |
| Conv4 output channels | 80 | [48, 64, 80, 96, 112, 128] | Filters in 4th convolutional layer |
| Conv1 kernel size | 11 | [3, 5, 7, 9, 11] | Kernel width in 1st convolutional layer |
| Conv2 kernel size | 5 | [3, 5, 7, 9] | Kernel width in 2nd convolutional layer |
| Conv3 kernel size | 7 | [3, 5, 7] | Kernel width in 3rd convolutional layer |
| Conv4 kernel size | 3 | [3, 5] | Kernel width in 4th convolutional layer |
| FC1 output features | 48 | [24, 32, 40, 48] | Neurons in 1st FC layer |
| FC2 output features | 48 | [24, 32, 40, 48] | Neurons in 2nd FC layer |
| Pooling type | Average | [Average, Max] | Pooling type between convolutional layers |
| Leaky ReLU slope | 0.055 | 0.001 to 0.1<br>(log-uniform) | Negative slope of leaky ReLU activation |
| Batch size | 20 | [8, 10, 12, 14, 16] | Samples per training batch |
| Random seed | 7 | [0, 7, 14, 21, 28, 35, 42] | Random seed for reproducibility |

### Threshold-crossing classification

For each z-scored t-series, we computed the standard deviation (*σ*) and a detection threshold of either 1.96 ∗ *σ* (containing 95% of data point of a standard normal distribution) or a more conservative 2.5 ∗ *σ* (containing > 98% of data point of a standard normal distribution). Standard deviation was calculated in a time window of 0.625 s prior to the stimulation onset. A time window of 0.375 s following the stimulation onset was defined as the detection range. Traces were classified as containing a calcium event if the signal exceeded the threshold for at least 2 consecutive time points within this window.

### Logistic regression classifier

The same labeled dataset used to train the deep neural network was used to train a logistic regression classifier that expresses probability of containing a calcium event as:

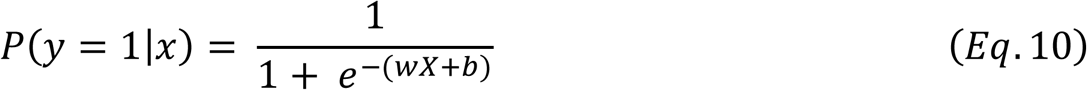

where *X* is the z-score time series, *w* is a weight vector, and *b* is the intercept. Data were split into training (80%) and validation (20%) sets. Hyperparameters, including the inverse regularization strength C and the L2 penalty, were optimized via grid search with 5-fold cross-validation. To account for label imbalance, class weights were adjusted to increase the penalty for errors in the minority class.

### Data analysis and representation, statistics and code availability

All packages, data analysis and graphical representations were written in Python using custom scripts and standard scientific libraries (*e.g. NumPy*, *SciPy*, *pandas*, *scikit-learn, pytorch*). Data are indicated throughout the text as mean ± standard error of the mean (SEM). Data are shown as individual points (gray circles) next to box plots. Red lines indicate the mean and SEM. Boxes represent the interquartile range (IQR) with the median marked by a horizontal line. Whiskers extend to data within 1.5 × IQR; points beyond this range are considered outliers.

Code is publicly accessible via a GitHub repository upon publication.

---

**Figure S1.**
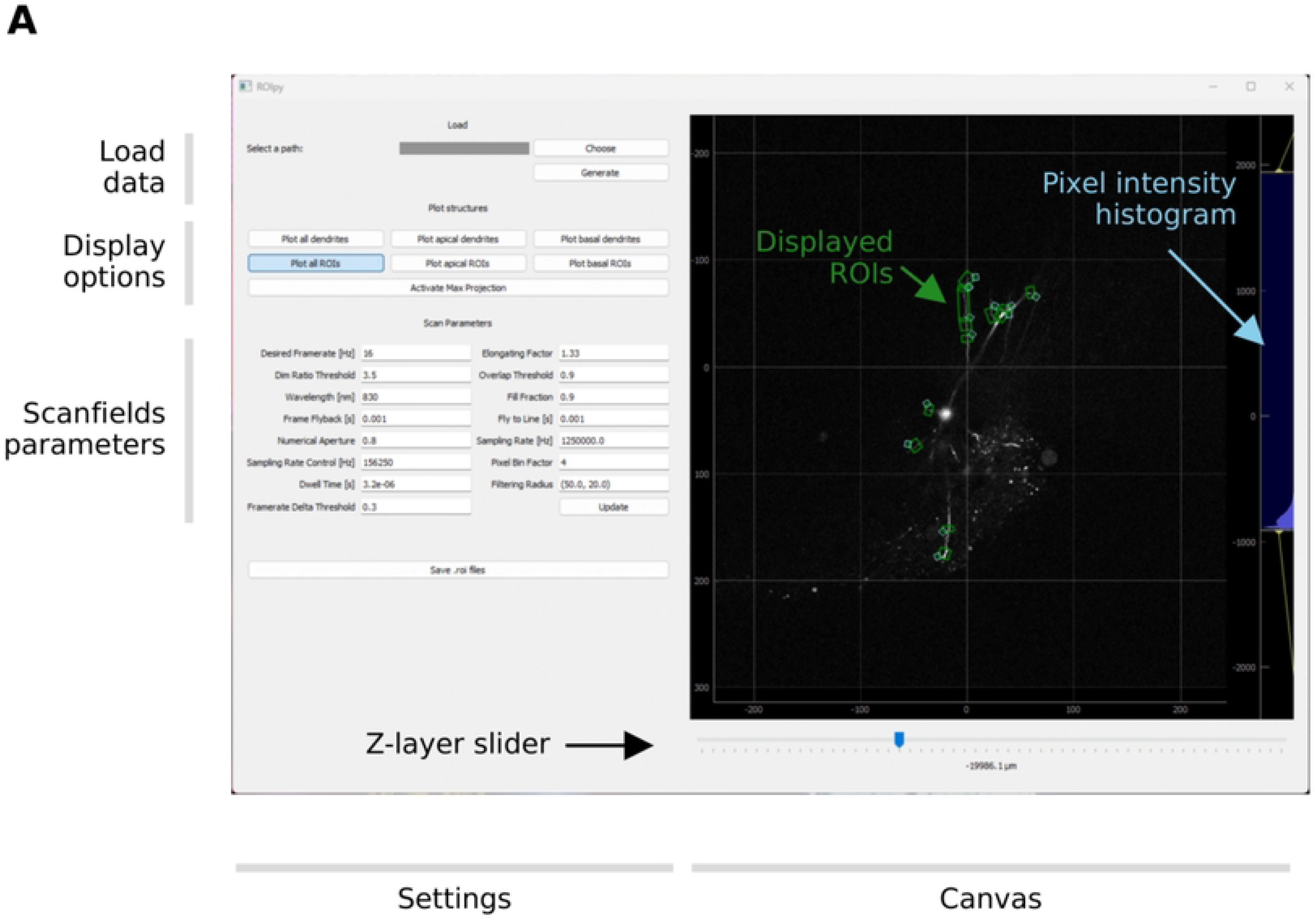
**A.** Graphical user interface (GUI) of *ROIpy*. The main layout consists of a setting panel and an interactive canvas. The settings panel includes options for loading data, interactive visualization controls such as toggling overlays, and adjusting scanfield parameters. The canvas provides a z-layers navigation slider and includes a tunable pixel intensity histogram (blue). Placed ROIs (green) are displayed in reference to the underlying dendritic morphology.

**Figure S2.**
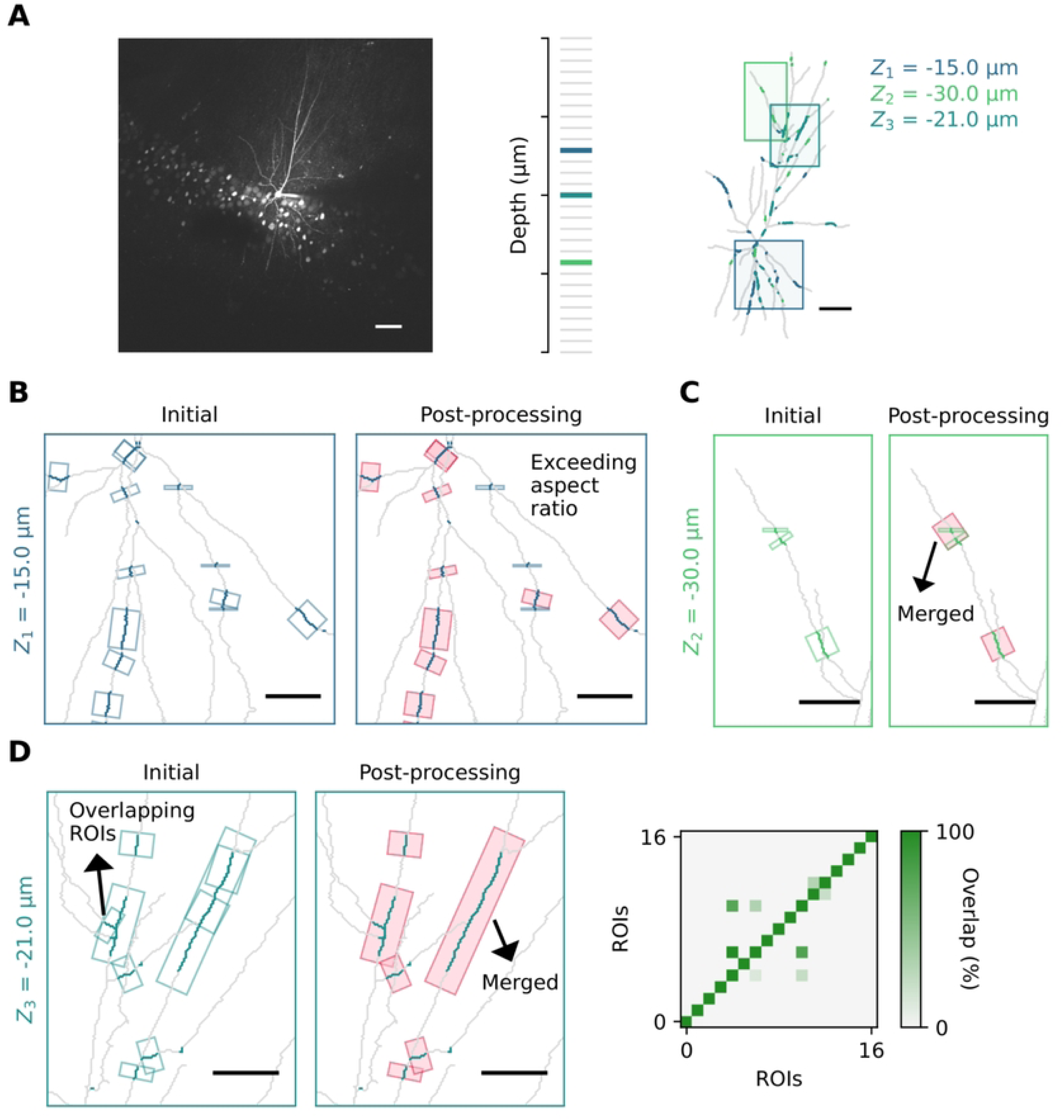
ROI generation and post-processing steps. **A.** Left: maximum intensity projection of a representative image stack showing a patched CA1 PN filled with A594. Scalebar: 50 µm. Center: three example z-planes extracted from the same stack. Right: traced neuron backbone, with dendritic segments color-coded by the z-plane wherein they are located (Z1 in blue, Z2 in green and Z3 in cerulean). Colored squares indicate the dendritic regions shown in panels B–D. **B.** Roi filtering based on aspect ratio. Left: initial ROIs placed on coplanar dendritic segments (blue). Right: retained ROIs after filtering (red); filtered ROIs exceeding aspect ratio threshold (blue). Scalebar: 20 µm. **C.** Merging of adjacent and coplanar ROIs. Left: pre-processing placement of ROIs on dendritic segments. Right: post-processing showing merged ROIs. Scalebar: 20 µm. **D.** ROI overlap filtering and merging. Left: example group with overlapping ROIs. Center: filtered and merged ROIs after processing; most overlapping ROI removed. Scalebar: 20 µm. Right: overlap matrix indicating the fraction of each ROI’s area covered by another ROI

**Figure S3.**
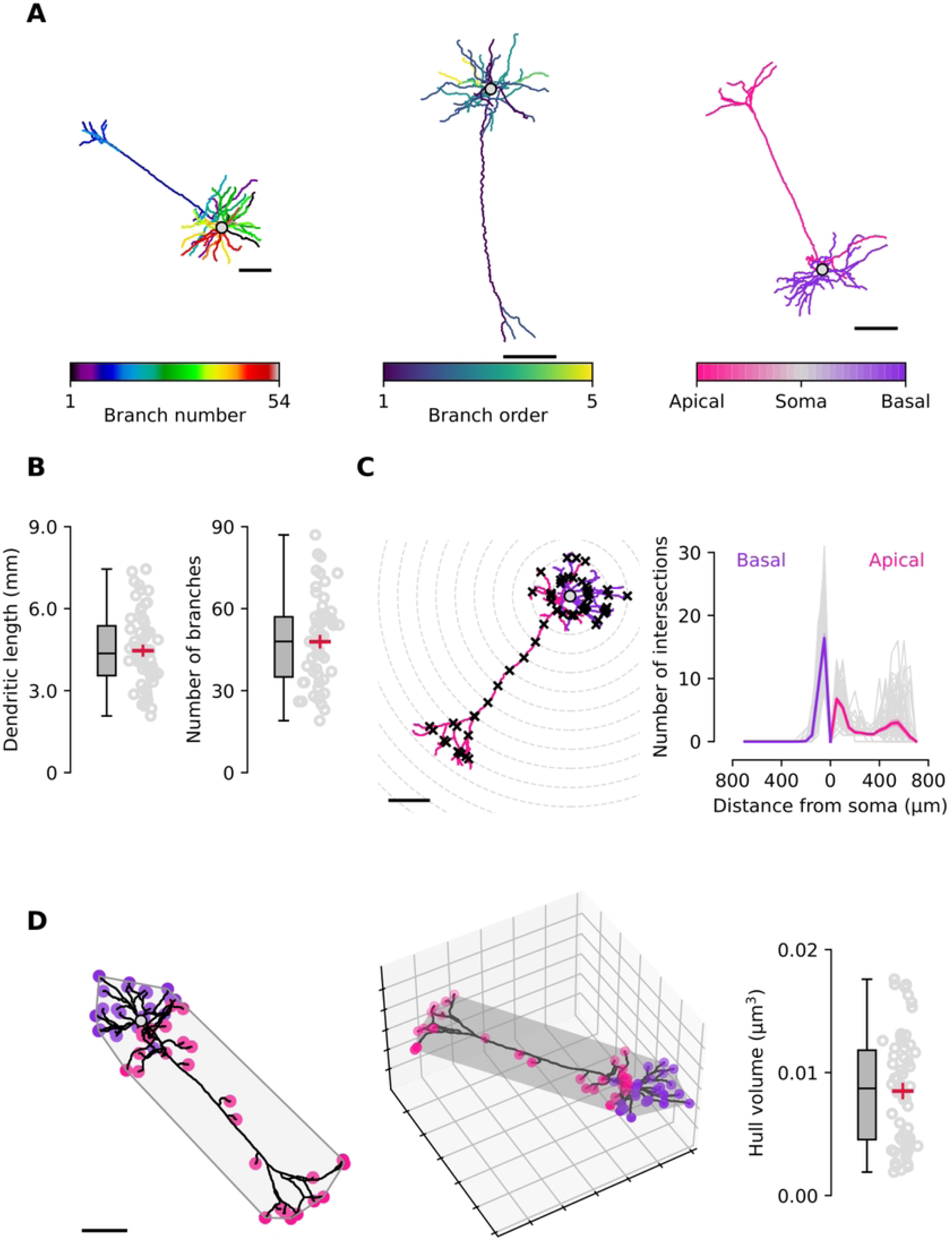
Morphological analysis of L5 pyramidal neurons using *ROIpy*. **A.** Representative morphological reconstruction of L5 PNs of the VISp, obtained from the Allen cell-type database and processed with *ROIpy*. Left: branches color-coded as individual segments from the soma or bifurcation points. Center: branches mapped by their branching order (*e.g.*, first order from soma, second order from first order, etc.). Right: branches grouped by dendritic compartment (apical vs. basal). Scale bars: 100 µm. **B.** Summary box and scatter plots for total dendritic length (left) and number of branches (right). **C.** Sholl analysis. Left: schematic of concentric circles (50 µm step) centered on the soma to quantify dendritic complexity. Right: number of dendritic intersections with Sholl circles as a function of radial distance, plotted separately for apical and basal branches. Individual neurons are shown in gray; average profiles with SEM shading are shown in pink (apical) and violet (basal). **D.** Convex hull volume analysis Left: 2D projection of the minimal convex hull enclosing dendritic endpoints. Scalebar: 100 µm. Center: 3D hull view with 100 µm grid spacing. Right: box and scatter plot of hull volume for every neuron in the dataset.

**Figure S4.**
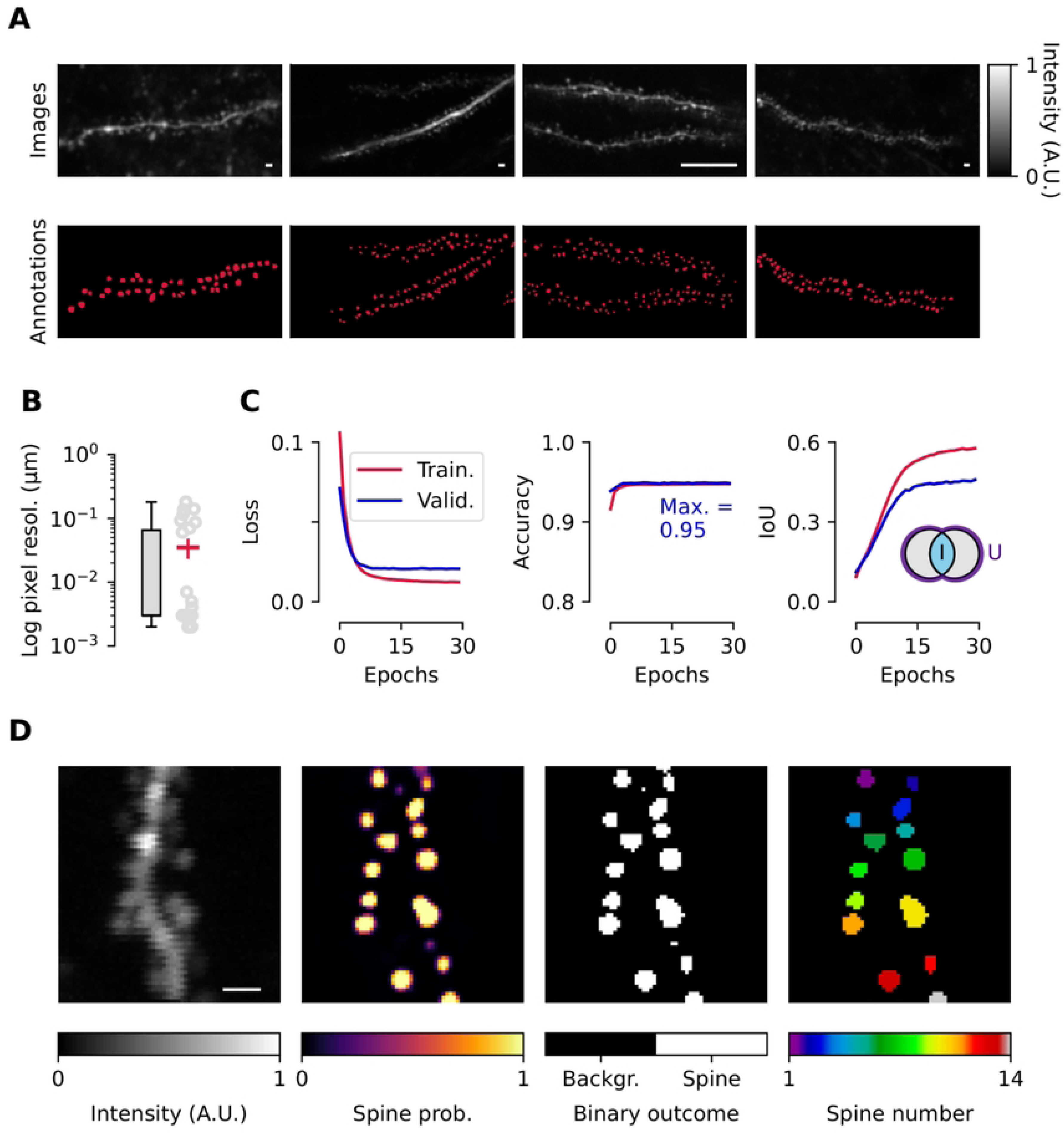
DeepD3 model training. **A.** Representative training set images. Top: raw images of A594-filled dendrites of vCA1 neurons. Bottom: the corresponding manual pixel-precise spine annotation (red). Scalebar: 2 µm in all images. **B.** Log distribution of pixel resolution of dataset images, reflecting variable acquisition settings across the dataset. **C.** Training performance. Left: training and validation loss converges over 30 training epochs. Center: training and validation accuracy scores increase and reach a plateau. Right: intersection-over-union (IoU) for spines during training, indicating improved prediction consistency. **D.** Spine inference process. From left to right: max projection of a single ROI, pixel-wise probability map, binarized and post-processed annotation, individual spine segmentation. Scalebar: 2 µm.

**Figure S5.**
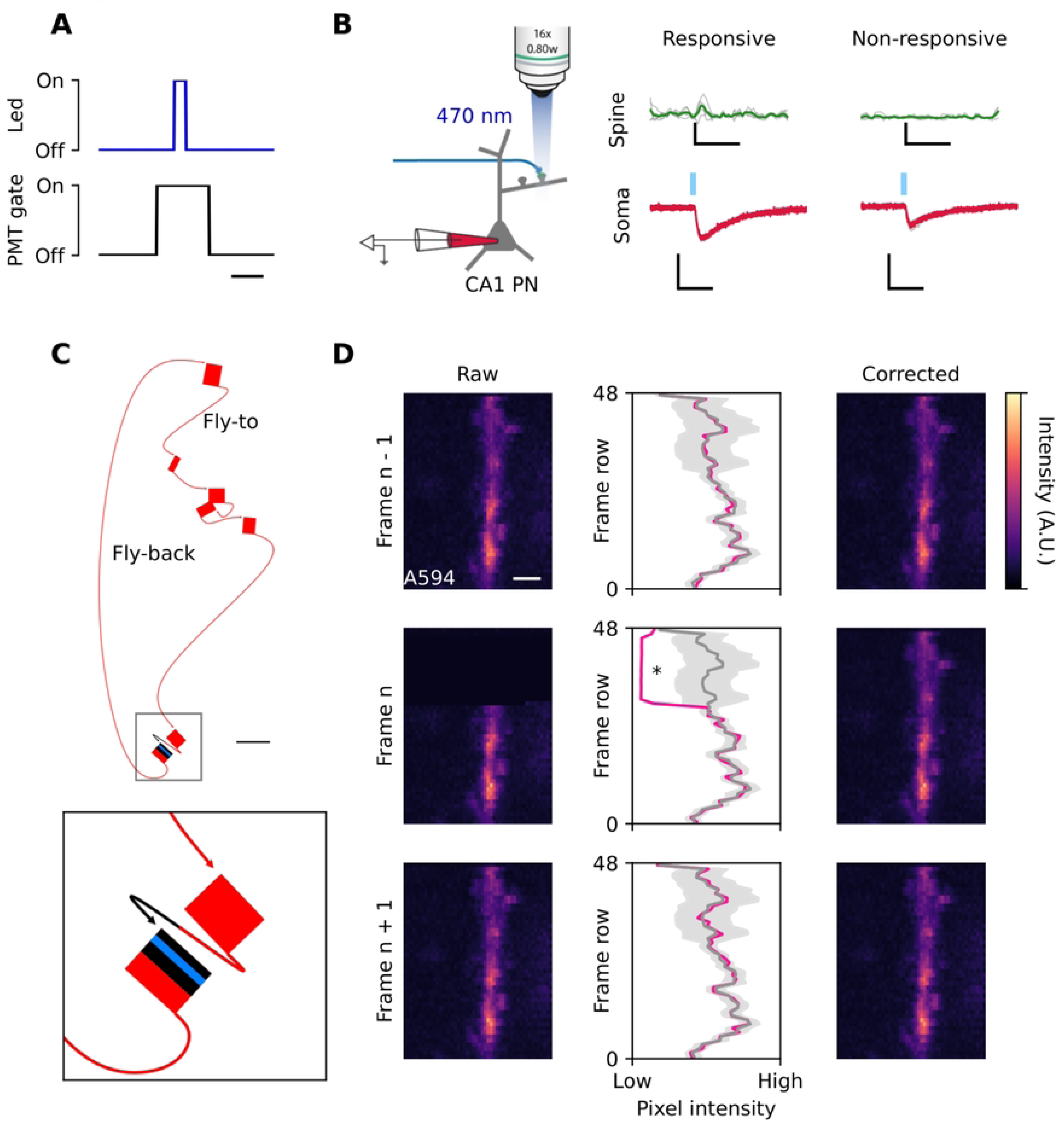
PMT gating and artifact correction during optogenetic stimulation. **A.** Digital-to-analog square functions showing timing relationship between blue-light delivery (blue trace, top) and PMT gate (black trace, bottom). The PMT gate is triggered shortly before light onset and remains active for 10 ms to prevent photodamage. This results in a horizontal dark artifact in the image (shown in panel D). Scalebar: 5 ms. **B.** Schematic (left) and representative data (right) showing ChR2-evoked activity in vCA1 PNs. Top traces depict calcium signals in a responsive and a non-responsive spine following light stimulation. Individual trials are plotted in gray and the average trace in green. Bottom trace shows the corresponding somatic EPSC recorded via whole-cell patch clamp. Individual trials are plotted in gray and the average trace in red. Calcium trace scalebars: 1 s, 0.5 ΔF/F_0_. EPSC traces scalebars: 100 ms, 50 pA. **C.** Top: schematic of the scanner path and PMT gating during single z-layer imaging. Red rectangles indicate scanned sparse ROIs; curved arrows represent connecting travel (fly-to next ROI and fly-back to start position). Bottom: zoom-in of the PMT gate (black) activation just before scanning the ROI (as indicated by the partially black colored arrow). During this inactive period, blue light illuminates the sample (light blue line). Scalebar: 20 µm. **D.** PMT gating artifact correction. Left column: raw frames from before (n – 1), during (n), and after (n + 1) stimulation showing a segment of A594-filled dendrite. Middle column: row-wise average intensity for each frame. Gray lines represent the mean row intensity across all 50 frames, with shaded areas indicating the tolerance threshold of 3*σ*. Pink traces correspond to the intensity profile of the current frame; rows exceeding the threshold are marked (*). Right column: corrected frame (n), where affected rows are interpolated using the corresponding rows from adjacent frames. Scalebar: 50 nm.

**Figure S6.**
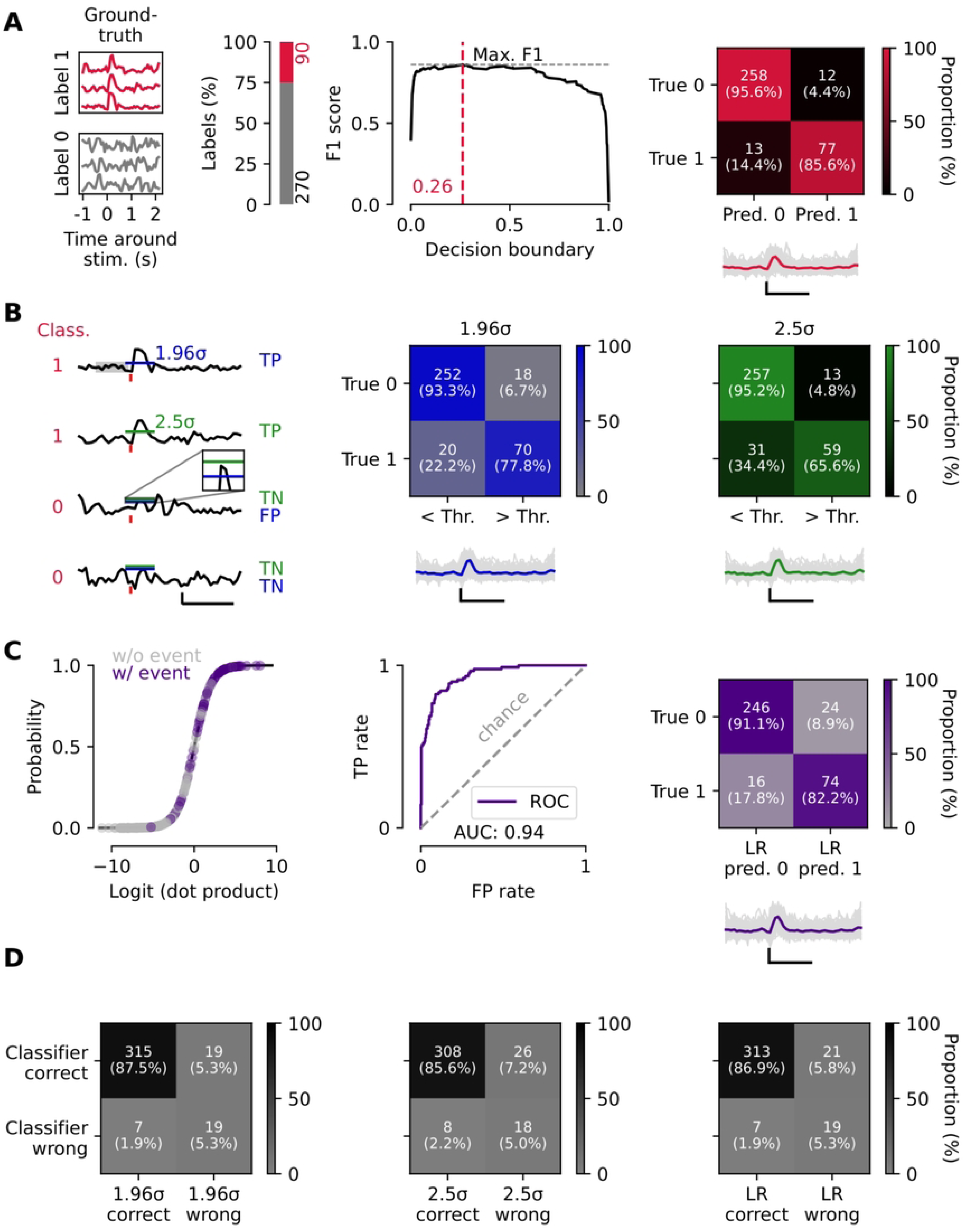
Classifier performance benchmarks and comparison. **A.** Model performance on test dataset. Left: representative ground truth z-scores for class 1 (red) and class 0 (gray) aligned to stimulation onset. Stacked bar plot indicates the class distribution of the test dataset. Center: F1 score as a function of the decision threshold on the test set. The maximum F1 value is indicated. Top-right: confusion matrix at the selected threshold, showing predicted vs. true labels. Bottom-right: detected z-scores are shown in gray and average in red. Scalebars: 1 s, 2 z-score units. **B.** Performance of threshold-based methods. Left: example traces with CNN model classification annotated in red. Stimulation is shown as a red vertical line. Gray shaded areas indicates the σ window. Blue and green horizontal lines indicate the threshold, with their length corresponding to the detection window. Blue and green annotations shown comparison with CNN classifier performance. TP: true-positive, TN: true-negative, FP: false-positive. Center and right: confusion matrices for detection using thresholds of 1.96 σ (left) and 2.5 σ (right). Thresholding at 2.5 σ reduces false positives but fails to detect most positive samples. Detected z-scores and average trace are shown below. Scalebars: 1 s, 2 z-score units. **C.** Logistic regression benchmark. Left: classifier output as a function of the logit (dot product of weights) is indicated with a black line. Overlayed dots are individual z-scores as classified by the regression model, colored by their ground truth annotation. Center: ROC curve with AUC. Top-right: confusion matrix using the decision boundary that maximizes the F1 score. Bottom-right: detected z-scores are shown in gray and average in violet. Scalebars: 1 s, 2 z-score units.

